# Comparative single-cell multiomic analysis reveals evolutionarily conserved and species-specific cellular mechanisms mediating natural retinal aging

**DOI:** 10.1101/2025.09.09.673285

**Authors:** Pin Lyu, Isabella Palazzo, Yang Jin, Leah J. Campbell, Clayton P. Santiago, Rogger P. Carmen-Orozco, Thanh Hoang, Jared A. Tangeman, Alexander D. Park, Shawn H. Yang, Jianming Shao, Rui Chen, David Hyde, Jiang Qian, Seth Blackshaw

## Abstract

Biological age is a major risk factor in the development of common degenerative retinal diseases such as age-related macular degeneration and glaucoma. To systematically characterize molecular mechanisms underlying retinal aging, we performed integrated single-cell RNA- and ATAC-Seq analyses of the retina and retinal pigment epithelium (RPE) across the natural lifespan in zebrafish, mice, and humans. By profiling gene expression and chromatin accessibility, we identified extensive cell type- and species-specific aging-dependent changes, with a much smaller number of broadly expressed and conserved genes that include regulators of inflammation and autophagy. We constructed predictive aging clocks for retinal cell types and observed dynamic, reversible shifts in cellular age following acute injury. Spatial transcriptomic analysis revealed region-specific aging signatures and proximity effects, with Müller glia exhibiting pro-rejuvenating influences on neighboring neurons. Targeted Müller glia-specific induction of Yamanaka factors reduced molecular age in rod photoreceptors and bipolar cells without altering glial age. Our findings define conserved and divergent regulatory and signaling pathways mediating retinal aging, highlighting Müller glia as potential therapeutic targets for combating age-associated retinal dystrophies.

## Introduction

Biological aging is a common phenomenon that is observed across almost all animal species, and is a major risk factor associated with the onset of a broad range of multifactorial disorders, ranging from cancer to neurodegenerative diseases. At the cellular level, aging is typically associated with a series of changes collectively referred to as hallmarks of aging, which include nuclear genome instability, mitochondrial dysfunction, formation of protein aggregates, cellular senescence, telomere shortening, DNA methylation, chronic inflammation, changes in intercellular communication, and other processes ^1^. Furthermore, identifying molecular aging clocks has made it possible to measure cellular age and the potential therapeutic effects of rejuvenative interventions ^2–4^. Identifying common genomic regulatory mechanisms that control cellular aging and developing therapies that slow or reverse this process hold tremendous potential in treating a wide range of age-related diseases ^5–7^.

However, analysis of the molecular mechanisms that underlie cellular aging has proved extremely challenging. By its very nature, aging is often a slow process, and aging-dependent molecular changes are relatively small and noisy, particularly for long-lived species such as humans ^1^. This stands in sharp contrast to the robust, temporally compressed processes such as embryonic development and response to acute injury ^8,9^. Hallmarks of aging often show strong cell type-dependent variation; for instance, telomere shortening is confined to actively mitotic cells and absent in tissues that are fully postmitotic, such as neuronal cells ^10^. Finally, it is becoming clear that age-dependent changes in gene expression show extensive tissue, region and species-specific variation ^11–17^. To best identify common mechanisms that mediate therapeutically-relevant aging-dependent changes in gene expression, gene regulation, and cell-cell signalling, there is a pressing need to generate robust age-matched single-cell multiomic atlases from organs and tissues that exhibit a high burden of age-related disease across both multiple cell types and species.

The retina represents an ideal system to perform these studies. Retinal anatomy and cell types are broadly conserved across all vertebrates, making it possible to directly compare aging-dependent changes in gene expression and regulation, both across species and cell types. Retinal aging affects many different aspects of vision ^18^ and is associated with characteristic changes in cellular morphology and function ^19–21^. Since the retina is an integral part of the central nervous system, insights gained from studying retinal aging are therefore likely to be more broadly relevant to normal and pathological CNS aging ^22–26^. Finally, biological age is a key risk factor for many blinding diseases, such as macular degeneration and glaucoma ^27,28^. Understanding the molecular details that are associated with aging at the single-cell level is critical to appreciate the onset and progression of these diseases.

In this study, we perform a comprehensive multiomic single-nucleus RNA (snRNA-Seq) and ATAC-Seq (snATAC-Seq) analysis over the course of natural aging, of both the retina and retinal pigment epithelium (RPE) in zebrafish, mice, and humans. This comprehensive analysis of aging-dependent changes in gene expression and chromatin accessibility was then used to identify differentially-expressed genes, generate cell type-specific aging clocks, and reconstruct aging-dependent gene regulatory and cell-cell signaling networks in all three species. Aging-dependent changes showed extensive cell type and species-specific variation, although a small number of broadly-occurring and/or evolutionarily-conserved changes in gene expression and cell-cell signaling were identified; notably these changes included age-dependent increases in inflammation and autophagy. Spatial transcriptomic analysis of aging-dependent changes in mouse retina validated the many changes observed using snRNA-Seq, identified distinct patterns of aging-regulated changes in central and peripheral retina, and identified retinal Muller glia as exerting a pro-rejuvenative effect on retinal neurons. This combined dataset represents a comprehensive view of aging-dependent changes across all retinal cell types, and identifies candidate mechanisms that mediate age-dependent cellular dysfunction associated with common retinal dystrophies.

## Results

### Generation of a single-cell multiomic atlas of aging zebrafish, mouse, and human retina

To comprehensively profile aging-dependent changes in gene expression and regulation at the cellular level, we performed combined snRNA- and ATAC-Seq at multiple timepoints across the full natural lifespan of zebrafish and mice. In zebrafish, we profiled retina and RPE from 1, 3, 6, 12, 18, 22, 24, 30, 36, and 48 months of age, with 8-10 samples pooled in each of 1-2 experimental replicates (Fig. 1A). For mice, we profiled retina and RPE from 5, 12, 17, 32, 49, 68, 91, 106, and 120 weeks of age analyzing a pool of 2 male and 2 females in 2 experimental replicates. Additional RPE enriched samples from mice were collected at 5, 49, and 91 weeks of age. Whole human scRNA-seq retina data from 104 healthy donors were obtained from the CZ CellXGene single-cell Portal ^29^, and ranged in age from 10 to 90 years of age, although samples from older donors (>70 years) predominated. Human RPE snRNA-seq samples and whole retina snRNA-Seq and scATAC-seq data were provided by Rui Chen’s group, and have been partially described elsewhere ^30^. This represents a total of 163,999, 152,796, and 3,079,521 retinal and RPE cells profiled from zebrafish, mouse, and human respectively.

**Figure 1:**
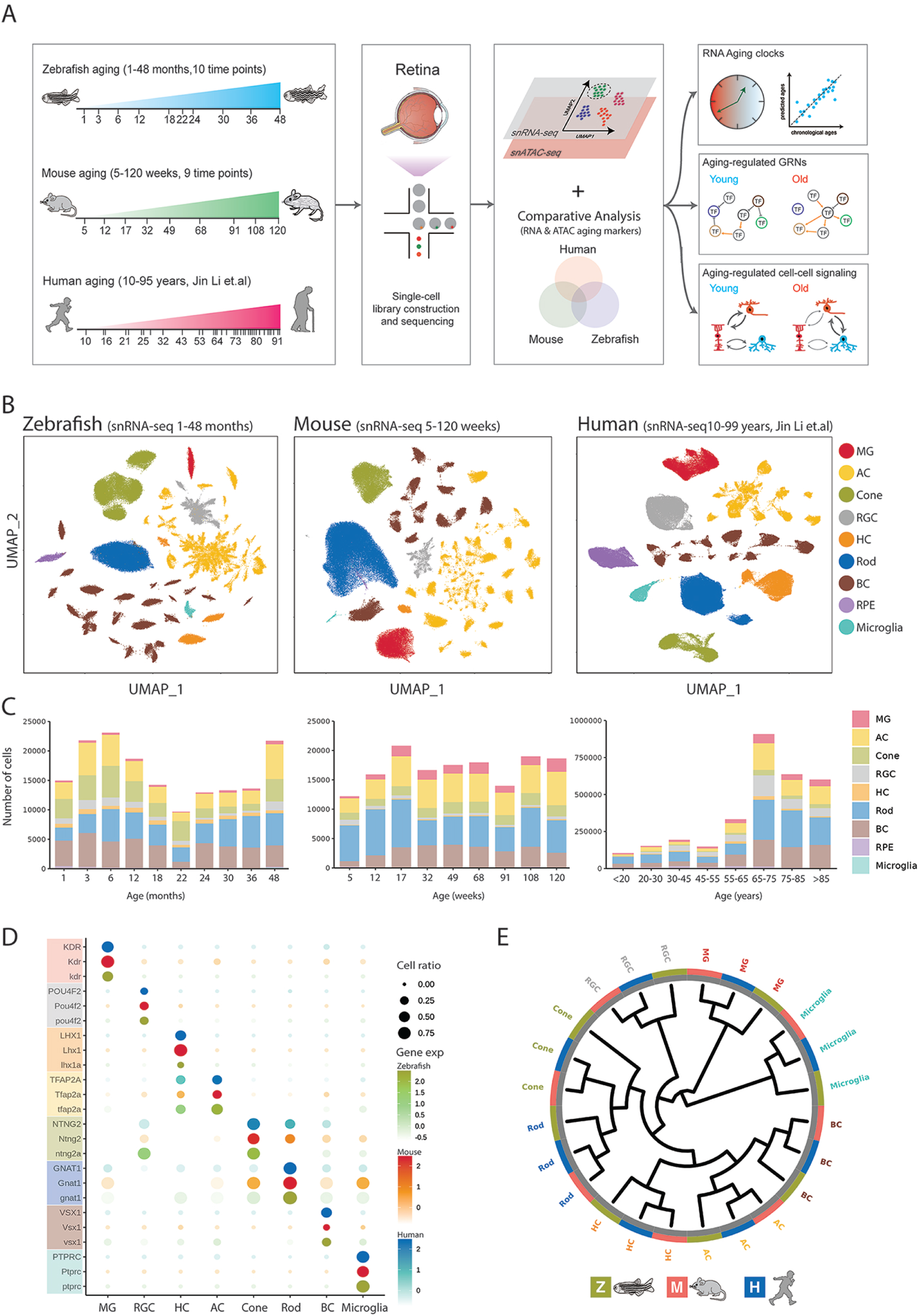
SnRNA-Seq analysis of naturally aging retina in zebrafish, mouse, and human. (**A**) Experimental design and computational workflow. Timelines depict age ranges and time points for all the samples in zebrafish (4–209 weeks), mouse (5–120 weeks), and human (10– 91 years, Jin et al.). Each sample undergoes snRNA-seq/scATAC-seq profiling, enabling high resolution comparative analyses of aging clocks, gene regulatory networks, and intercellular signaling across species. (**B**) UMAP embeddings of integrated single-cell transcriptomes for each species. Cells are color-coded by annotated major cell classes: Müller glia (MG), amacrine cells (AC), cones, retinal ganglion cells (RGC), horizontal cells (HC), rods, bipolar cells (BC), retinal pigment epithelium (RPE), microglia, and astrocytes (human only). (**C**) Cell-type composition across different ages. Stacked bar plots illustrate cell counts per major class at each time point for zebrafish, mouse, and human. The x-axis denotes chronological age, and the y-axis indicates absolute cell counts. (**D**) Conserved, cell-type-specific marker genes across the three species. Dot plots show average expression levels (color scale) and the fraction of cells expressing each marker (dot size) in zebrafish, mouse, and human. Selected transcription factors and canonical markers are listed on the y-axis. (**E**) Dendrogram plot depicting transcriptional homology among retinal cell types across species. Branch lengths reflect global gene-expression distances; ring colors indicate species. Cell types are labeled around the plot.

Clustering and UMAP analysis of both snRNA-Seq and snATAC-Seq from each species showed good resolution of all major cell types, including Muller glia (MG), rod photoreceptors, cone photoreceptors, bipolar cells (BC), amacrine cells (AC), retinal ganglion cells (RGC), horizontal cells (HC), microglia, and RPE (Fig. 1B, Fig. S1A, Table S1). This analysis also efficiently resolved retinal subtypes, with 23 bipolar cell subtypes detected in zebrafish, 15 in mice, and 12 in humans, corresponding precisely to previously reported numbers ^31–33^. The relative fraction of major retinal cell types remained largely consistent across the lifespan in all three species, and gene expression and chromatin accessibility were associated with known cell type-specific markers in all cases (Fig. 1C-D, Fig. S1B-C). Annotated cell types from different species showed similar gene expression and chromatin accessibility profiles, with mouse and human consistently more similar to one another than either were to zebrafish (Fig. 1E, Fig. S1D). This demonstrates that the datasets obtained here are of high quality, and efficiently profile all major retinal and RPE cell types across the lifespan.

### Aging-dependent changes in gene expression in zebrafish, mouse, and human

To identify age-dependent gene expression changes across all major retinal and RPE cell types, we applied GAM fitting across the lifespan for each gene in each cell type and species, followed by k-means clustering across aging time points to identify major patterns of change (Fig. 2A). In line with studies of human cellular aging in other tissues ^34–36^, we observe substantial sex-dependent differences in the aging human retina (Fig. S6, Table S5). We identified 1,805 genes enriched in young individuals (<40 years old) and 3,118 genes enriched in aged individuals (>70 years old) across both sexes. Additionally, 364 genes were enriched in young females and aged males, and 187 genes in aged females and young males, indicating sex-dependent expression patterns. To avoid potentially confounding effects of sex, we therefore initially analyzed male and female human samples separately. Our analysis identified 10-12 age-dependent gene clusters for each species, which further grouped into three categories: enriched in young samples, enriched in aged samples, and either up- or down-regulated in middle age (Fig. 2B-E). This yielded between 1,153 and 5,059 aging-associated genes in each cell type (Fig. 2B).

**Figure 2:**
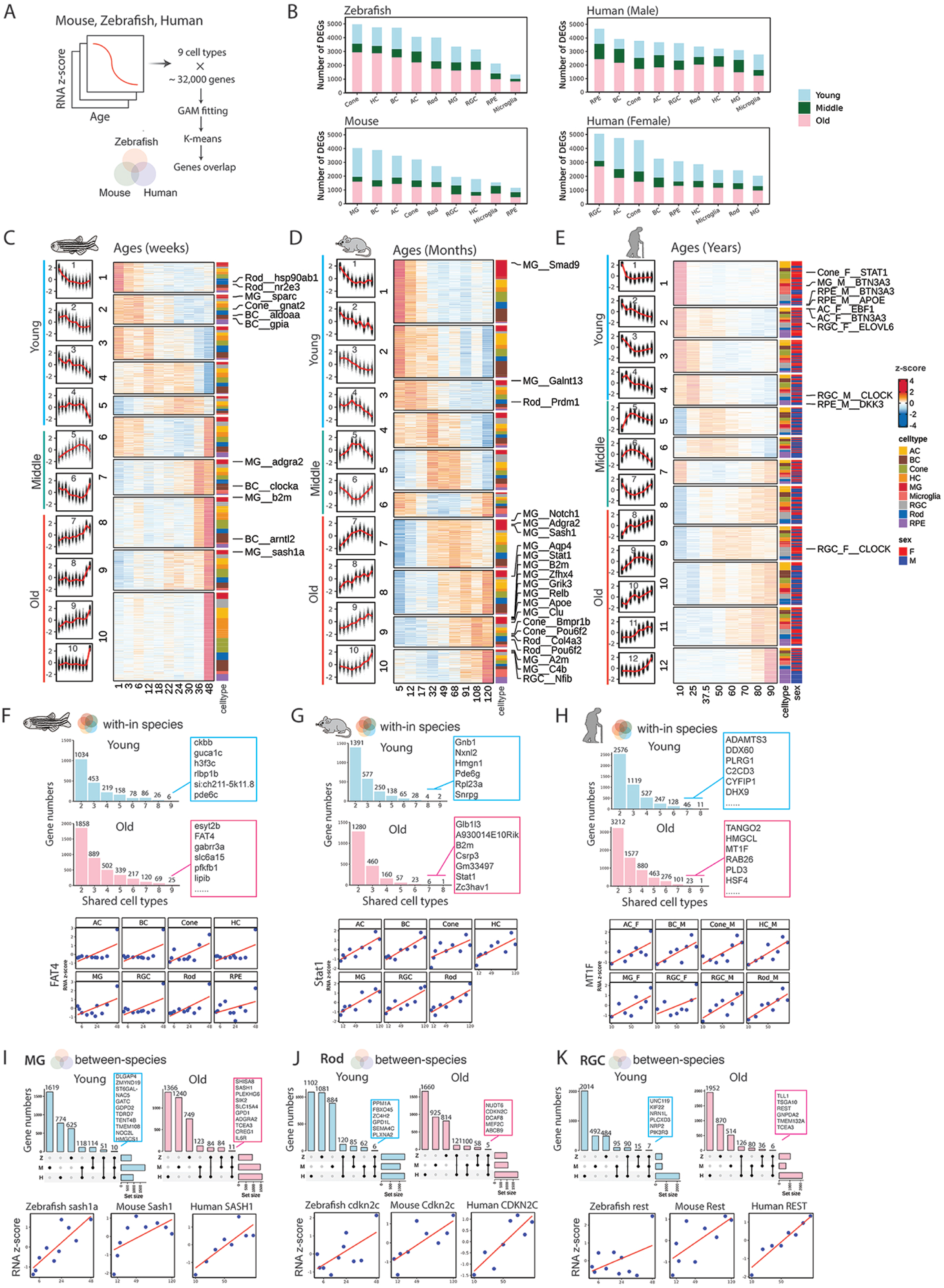
Aging-regulated genes identified in zebrafish, mouse, and human retina. **(A)** Computational pipeline for identifying aging-associated genes. For each species (zebrafish, mouse, human), we fit generalized additive models (GAMs) to cell type-specific RNA Z-scores over age (covering ∼18,000 genes x 9 major cell classes), clustering trajectories of aging-regulated genes by k-means, and comparing clusters across species to identify aging-regulated DEGs. (**B**) Counts of aging-associated genes in old vs. young for each cell type and species. Bars above zero (pink) denote genes increased with age; bars below zero (blue) denote genes decreased with age. These bar plots are separated by zebrafish (top), mouse (middle), and human (male and female). (**C-E**) Heatmaps showing the distinct clusters of aging-regulated genes in (**C**) zebrafish, (**D**) mouse, and (**E**) human. Within each species, clusters are ordered by overall trend (Young, Old and middle aged). Representative aging associated genes from Müller glia (MG), rods, cones, RPE, etc., are annotated on the right of each heatmap. The left scaler plot shows the overall gene expression pattern for each cluster. (**F-H**) Within-species overlaps of aging-associated genes among different cell types. Bar plots demonstrate the number of aging-associated genes shared among multiple cell types (> 1 cell types shown) for each species (zebrafish, mouse, human). Below each histogram, example trajectory plots for a selected genes including *fat4* in zebrafish (**F**), *Stat1* in mouse (**G**), and *MT1F* in human (**H**) illustrate consistent changing tendency across multiple cell types. (I-K) Cross-species overlaps of conserved aging genes in three representative cell types: (**I)** Müller glia, (**J**) rods, and (**K**) retinal ganglion cells (RGC). UpSet plots (top) depict the number of shared increasing (left) or decreasing (right) genes across zebrafish (z), mouse (m), and human (h). Bar plots (bottom) show trajectory examples for one conserved gene per cell type in each species (e.g., *SASH1* in MG, *CDKN2C* in rods, *REST* in RGC).

In zebrafish and in human females, we consistently observed more aged-enriched genes across all cell types, while in moice, smaller numbers of young and aged-enriched genes were generally observed. In all three species, genes that were dynamically expressed in middle age generally remained a relatively small fraction of the total number of age-regulated genes (Fig. 2B). Levels of age-dependent variation in gene expression differed considerably by cell type, species, and sex. In zebrafish, however, the cell types that showed the highest number of aging-regulated genes were cones, HC, and BC, while in mice MG showed the highest number. In humans (leaving aside RPE samples where donors skewed heavily male), female samples showed the highest levels of age-dependent variation in RGCs, but in males RGCs showed lower levels of age-dependent variation (Fig. 2B). A full list of all aging-regulated genes in each species is shown in Table S2.

In each species we observe 4 clusters of genes enriched in young samples and 4-5 clusters enriched in aged samples, which differ both in the kinetics and magnitude of changes in gene expression (Fig. 2C-E). The great majority of observed aging-regulated changes in gene expression were both cell type and species-specific, as were most of the functional categories identified by Gene Ontology (GO) analysis (Table S3). However, there were a limited number of common functional categories that were shared across cell types and often across species, although categories increased with age were generally associated with established hallmarks of aging. Young cells consistently expressed higher levels of genes regulating translation and RNA processing, as well as many cytoskeletal components. In contrast, aged cells consistently showed higher expression of genes related to autophagy, inflammation, cell adhesion and morphology, intracellular transport, and response to cellular stress. Specific cell types exhibited stronger functional signatures, with inflammatory signaling particularly strong in MG and microglia, autophagy and ciliary-related genes strong in rod and cone photoreceptors and RGCs (Fig. S4). Genes specific to resting MG such as *Aqp4*, *Glul*, and *Apoe* along with Notch pathway components were upregulated with age in mice, which also showed the strongest inflammatory gene signatures in aged MG, potentially reflecting a compensatory anti-inflammatory response (Fig. 2D, Table S2) ^37,38^.

In many tissues, aging leads to the accumulation of senescent cells. These cells undergo replicative arrest, secrete pro-inflammatory factors, and express a characteristic set of genes. We investigated whether genes enriched in aged cells were enriched for senescence-associated genes in any of the species profiled. No cell types in aged zebrafish were enriched for senescent cells, using either the SenMayo or Reactome senescence gene list ^39,40^ (Fig. S5, Table S4). In contrast, mouse Muller glia and microglia were strongly enriched for SenMayo senescence-associated genes, reflecting the generally stronger age-dependent induction of inflammatory gene expression in mice. In humans, SenMayo and Reactome-associated genes were both enriched in Rod and RPE, whereas SenMayo genes were specifically enriched in aged microglia. Overall, age-dependent expression of senescence-association genes was modest and highly cell type and species-specific.

While the great majority of aging-regulated genes in all three species showed altered expression in only two cell types or less, in each species a small number of genes that did show common aging-dependent changes in expression in 8 cell types or more were identified (Fig. 2F-H, Table S2). In some cases, these include highly cell type-specific genes such as the cone-specific *guca1c* and *pde6c* in zebrafish and the rod-specific *Pde6g* in mice, and therefore likely reflected ambient contamination. However, in most cases these genes that were indeed broadly expressed and fell into common aging-regulated functional categories identified using GO analysis. These include the cell adhesion factor *fat4* in zebrafish; the inflammation-associated genes such as *B2m* and *Stat1*, the ribosomal protein *Rpl23a*, and the splicing factor *Snrpg* in mice; and stress response factors such as *MT1F* in humans.

While very few individual genes showed consistent aging-regulated changes in expression across many cell types in all species, a number were identified for each major cell type. In aged MG, for instance, we observe increased expression of the inflammatory-associated factors *Sash1* and *Sik2* and the cell adhesion GPCR *Adgra2* in all three species. In rod photoreceptors, we observe consistent upregulation of the proinflammatory factor *Ndut6* and the cyclin-dependent kinase inhibitor *Cdkn2c*, which has been implicated in cellular senescence^41^. RGCs upregulate the inflammation-associated genes *Tll1* and *Tsga10*, along with the transcription factor *Rest*, which represses expression of neuronal genes (Fig. 2I-K).

### Histological and physiological analysis of aging-related changes in zebrafish and mice

To validate predicted age-dependent changes in inflammatory signaling and cell morphology, and to better interpret the functional significance of observed aging-dependent changes in gene expression, we performed longitudinal histological analysis across the normal lifespan of both zebrafish and mice. In the zebrafish retina, we observe a progressive retraction and shift to a helical, rather than rod-like, structure of the MG basal processes (Fig. S2A, A’).

This correlates with a progressive thinning of the inner plexiform layer (Fig. S2B), which fits with the previously described thinning of the aging zebrafish retina ^21^. We also observed an increase in the number of Lcp1-positive microglia and macrophages (Fig. S2C, D).

To validate age-dependent changes in expression observed in zebrafish using snRNA-Seq, we also performed single molecule fluorescence *in situ* hybridization analysis for *sparc* and *dkk1b*, which respectively show decreased and increased expression over time in MG (Fig. S2E). As expected, we observe a clear decrease in *sparc* expression in MG throughout the retina (Fig. S2F, F’). Because the Sparc protein is involved in the formation and maintenance of the extracellular matrix ^42^, the decreased *sparc* expression in older fish could result in the loss of MG interaction with the basal inner limiting membrane. In contrast, the increased *dkk1b* expression with age was restricted to MG in the ventral retinal periphery (Fig. S2G, G’, G’’).

In mice, we observe a strong age-dependent increase in GFAP immunostaining, with basal processes of MG becoming strongly immunopositive by 84 weeks (Fig. S3A, B). Iba1, which stains microglia and macrophages ^43^, showed progressively increased signal in the outer nuclear layer (Fig. S3C, D). We observed an age-dependent increase in expression of the transcription factor Cux2 in our snRNA-Seq data, and confirmed this using immunostaining (Fig. S3E, F). We also took this opportunity to validate previously described age-dependent molecular and physiological changes in the mouse retina. We confirmed previously reported age-dependent intrusion of PKCalpha-positive apical dendrites of rod bipolar cells into the outer nuclear layer ^44,45^ (Fig. S3G,H). Finally, using electroretinogram (ERG) analysis, we also confirmed age-dependent decreases in the scotopic a-wave and both the scotopic and photopic b-waves, along with a decline in flicker response that was more prominent at 10 Hz than 30 Hz ^46^ (Fig. S3I-L). In conclusion, we observe a pervasive gliotic response in aged retinas and largely confirm previously-reported molecular and physiological age-dependent changes.

### Calculating snRNA-Seq-based aging clocks for retinal cell types

Previous studies have used both bulk and single-cell transcriptomic data to calculate cellular age in the brain ^4,47–49^, but this data is lacking for retina. We applied an elastic-net regression method ^50^ to generate cell type-specific aging clocks for the major retinal cell types of all three species. For each sample and cell type, we first applied KNN smoothing ^51^ to reduce noise for each single cell. Then we selected highly cell-type-specific aging-correlated gene features for aging clocks using Spearman correlation (detailed in supplementary methods), and supplement these features with genes previously implicated in mammalian aging obtained from AgingMap ^52^ and SenCID ^53^. Finally, with de-noised expression profiles and selected gene features, we built cell type-specific aging clocks for each species, with 70% sampled cells in each sample to train the model, and evaluated these models with the remaining 30% of cells (Fig. 3A).

**Figure 3:**
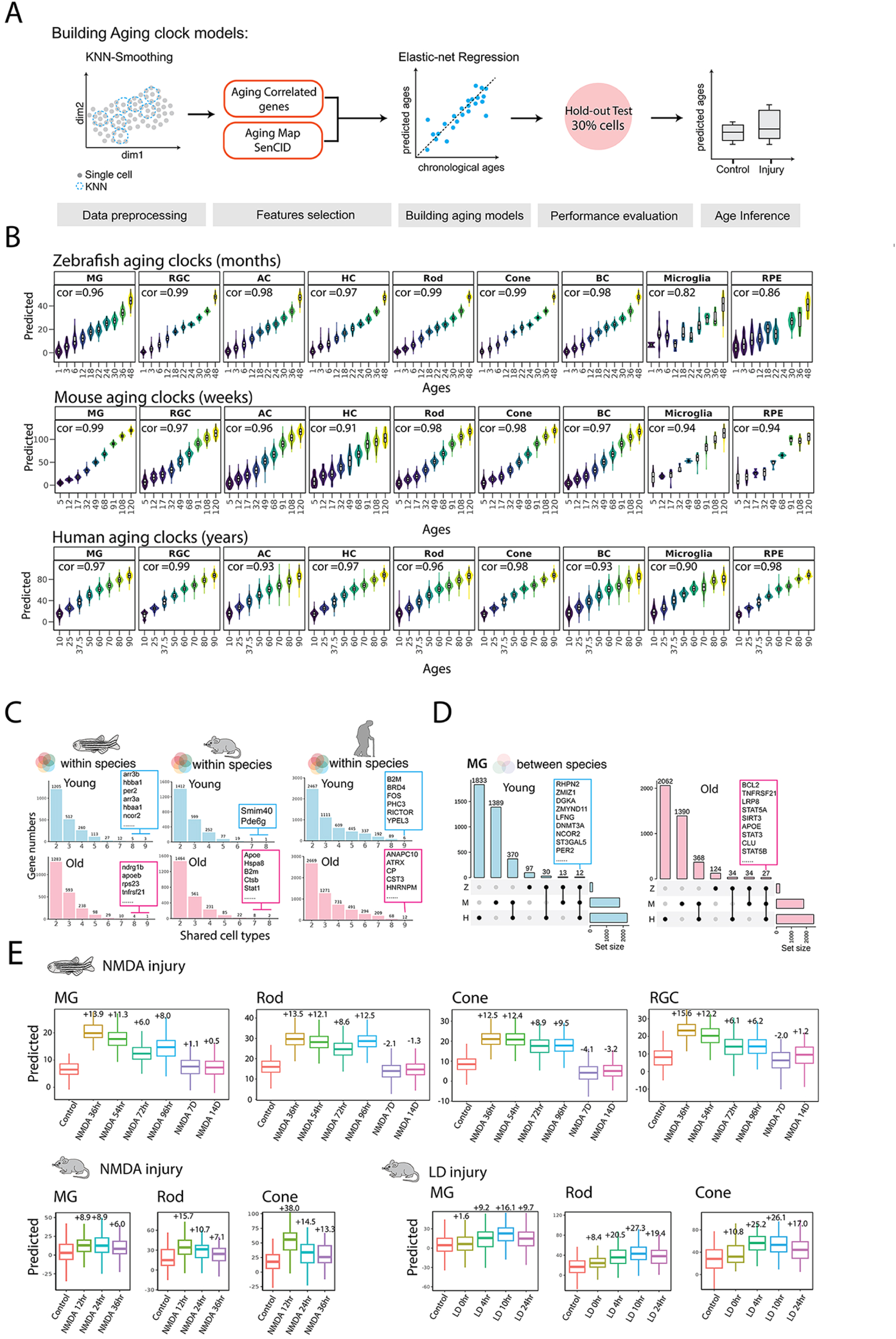
Cell type-specific aging clocks for major retinal cell types in zebrafish, mouse, and human retina. (**A**) Workflow for constructing cell-type-specific aging clocks. Transcriptomes are initially smoothed and denoised using KNN. Aging-correlated genes from the datasets are selected and combined with Aging Map and SenCID genes. Elastic-net regression models are trained to predict chronological age, validated on a 30% held-out subset and an independent Xenium spatial transcriptomic dataset, and subsequently applied to estimate ages in retina injury samples. (**B**) Predicted versus true ages for eight retinal cell classes in zebrafish (months), mouse (weeks), and human (years). Violin and box plots illustrate the distribution of predicted ages across true ages, with Pearson correlation coefficients (cor) provided in the upper left of each panel. (**C**) Application of mouse aging clock models to Xenium spatial transcriptomics data. Box plots depict predicted ages (y-axis) for each cell type (MG, RGC, AC, HC, rod, cone, BC, RPE) against the actual sample ages (x-axis), highlighting correlation (cor) indicating spatial profile concordance. (**D**) Within-species overlap of aging clock genes across different cell types. Bar plots demonstrate the number of clock genes shared among multiple cell types (> 4 cell types shown) for each species (zebrafish, mouse, human), with the top shared genes labeled. (**E**) Cross-species overlap of Müller glia (MG) aging clock genes, differentiated by up-regulated (right) and down-regulated (left) features using an UpSet plot. Horizontal bars indicate total feature set sizes per species (Z = zebrafish, M = mouse, H = human); vertical bars depict intersection sizes for each species combination. (**F**) Top panel showing predicted biological age in zebrafish retinal cells following NMDA-induced injury. Box plots illustrate inferred ages for MG, rod, and cone cells across control and six post-injury timepoints (36 hr, 54 hr, 72 hr, 96 hr, 7 d, 14 d), revealing transient aging responses. Bottom panel showing predicted biological age in Mouse retinal cells following NMDA and LD-induced injury. Box plots illustrate inferred ages for MG, rod, and cone cells across control and post-injury timepoints (0hr, 4hr, 10hr, 12hr,24hr, and 36hr), revealing transient aging responses. The median predicted age differences between injury and control are labeled on each boxplot.

We obtained 27 aging clocks after we applied the approach to each cell type in the three species. The number of genes in each clock ranged from 200 (HC in Zebrafish) to 4,001 (AC in Human Female) (Table S6). Each aging clock demonstrated a strong prediction of the biological age, with correlation coefficient of predicted and actual ages ranging from 0.86-0.99 (Fig. 3B).

Reflecting the properties of the snRNA-Seq data, the genes comprising each cellular aging clock were generally highly cell and species-specific, with several notable exceptions. *Apoe*, which is one of the genes most closely genetically linked to lifespan in humans ^54,55^, was shared across at least 8 cell types in zebrafish and mouse, and was aging-associated in MG in all three species. Other inflammation-associated genes such as *Tnfrsf21*, *B2m*, *Stat1,* and *Stat5a*, as well as stress-associated genes such as *Cp*, *Cst3*, *Cstb*, and *Hspa8* were also shared among large numbers of cell types and/or shared across species (Fig. 3C,D).

We next used the cell-specific clocks to assess the molecular age of different cell types following an acute retinal damage in zebrafish, which exhibit a MG-dependent regeneration of retinal neurons ^56,57^. Intravitreal injection of NMDA induces the loss of primarily retinal ganglion and amacrine cells, with a smaller percentage of photoreceptors and bipolar cells being lost ^58^, and leads to neurogenic reprogramming of MG, which then produces neuronal progenitor cells that go on to fully replace dying cells. Six-month old fish were treated acutely with NMDA and profiled using snRNA-Seq at multiple timepoints following injection, and cellular ages were estimated using our cellular aging clocks. We consistently find that measured cellular ages of MG, rod, and cone photoreceptors increase substantially within hours following injury, but return to baseline as the neurons are regenerated within 7 days (Fig. 3E). Finally, we applied our mouse aging clocks to previously published scRNA-Seq obtained from NMDA-treated retinas ^9^. As with zebrafish, we see that cellular age is significantly increased within 12 hours of NMDA treatment, eventually returning to baseline levels (Fig. 3E).

### Identifying aging-dependent changes in gene regulation

Disrupted transcriptional and epigenetic regulation is a known hallmark of aging, with loss of DNA methylation and broad increases in genome accessibility reported in some cases ^59–61^. To determine if this was the case in retinal cell aging, we analyzed our snATAC-Seq for age-dependent changes in chromatin accessibility. Here too, we observed dramatic cell type and species-dependent variation (Fig. S7, Table S8). In zebrafish, retinal ganglion and amacrine cells showed few age-dependent changes in the number of differentially accessible regions (DARs), and while the largest number of age-dependent DARs was seen in Muller glia and cones. Mice showed substantially fewer age-dependent DARs, and these were concentrated in rod and cone photoreceptors and Muller glia. Humans, in contrast, showed large numbers of aging-dependent DARs in each cell type, with aged cells consistently showing increased numbers of DARs.

Next, to identify changes in gene regulation that underlie transcriptional changes associated with cellular aging, we constructed the gene regulatory networks (GRNs) in each cell type by integrating scRNA- and ATAC-seq data from each species. By directly correlating patterns of transcription factor expression and footprinting-inferred binding at regulatory elements associated with target genes, we then identified both positive and negative age-dependent regulons, as previously described ^58,62^(Fig. 4A). We again observed substantial cell type and species-dependent variability in the number of age-dependent regulons, with zebrafish Muller glia, mouse RGC, and both zebrafish and mouse horizontal cells showing notably smaller numbers than other retinal cell types (Fig. 4B).

**Figure 4:**
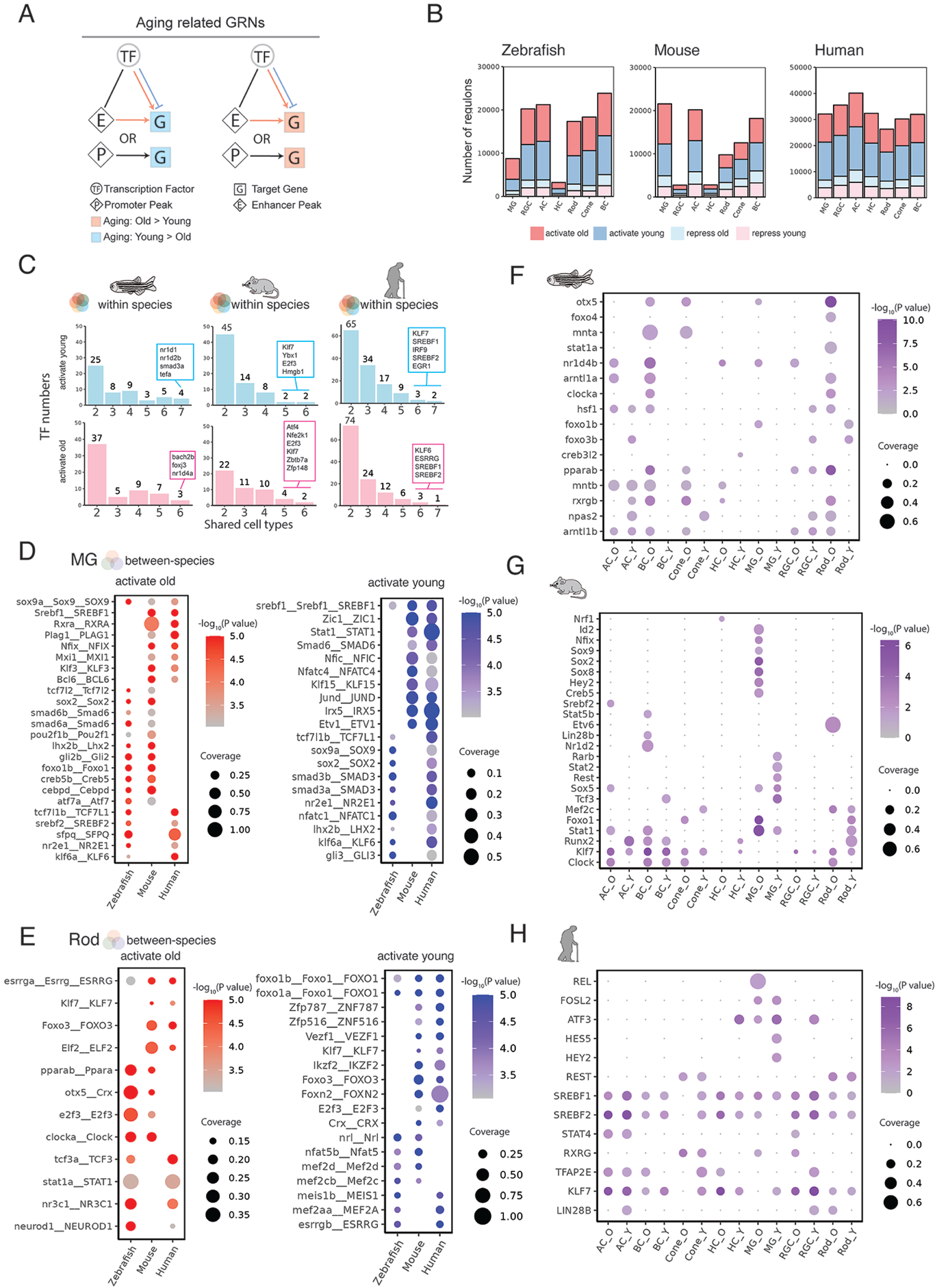
Transcription factors showing age-dependent changes in activity in zebrafish, mouse, and human retina. (**A**) Schematic overview of the strategy used to define aging-related GRNs. Transcription factors (TFs) are linked to target genes (G) via promoter (P) or enhancer (E) peaks. Differentially genes increasing or decreasing along ages (increasing: Old > Young, decreasing: Young > Old) are used to infer four classes of TF–gene regulatory relationships: Activation Old, Activation Young, Repression Old, and Repression Young. (**B**) Stacked bar plots showing the total number of regulons of each class in 7 major retinal cell types from zebrafish, mouse, and human. For each species, bars are grouped by cell type (MG, RGC, AC, HC, Rod, Cone, BC), and are colored by different regulatory classes. (**C**) Bar plots of within-species overlap of enriched transcription factors across retinal cell types separated by distinct regulatory categories (light blue: activation in young, pink: activation in old). For each species (zebrafish, mouse, human), bars indicate the count of TFs enriched in more than one cell type, and the most frequently shared TFs are labeled above their respective bars. (**D**) Dot-plot showing the shared TF enrichment (at least in 2 species) in Müller glia (MG) cells across zebrafish (Z), mouse (M), and human (H) in 2 regulatory categories (activate young and activate old). Dot size represents the proportion of target genes covered by each TF (coverage), and color represents the enrichment score (–log_10_P-value). (**E**) Dot-plot showing the shared TF enrichment (at least in 2 species) in Rod cells across zebrafish (Z), mouse (M), and human (H) in distinct regulatory categories (activate young and activate old). Dot size represents the proportion of target genes covered by each TF (coverage), and color represents the enrichment score (–log_10_P-value). (F-H) Dot-plot of transcription factor (TF) enrichment in zebrafish (**F**), mouse (**G**), and human (**H**) retinal cells across two regulatory categories: activation in young (_Y) and activation in old (_O). The x-axis labels each cell-type–regulatory category combination (suffix “_Y” = activate young; “_O” = activate old). Dot size denotes the proportion of that TF’s target genes covered (coverage), and dot color encodes the enrichment score (–log_10_P-value).

For each cell type, we obtained thousands of aging-related regulons, which in turn can be classified into four groups based on whether they activate or repress expression of young- or aged-enriched genes. The great majority of these regulons selectively activated expression of either young or aged-specific genes, and we focused our attention on these (Fig. 4B, Table S9). To identify key evolutionarily conserved regulatory nodes controlling cellular aging, we next tested each transcription factor to determine whether it was significantly enriched in genes activated in either young or aged cells across all three species. As with the snRNA-Seq analysis, most TFs were highly cell type and species-specific in their activity, and very few were shared across all species in any cell type. A number of genes, however, did show evolutionary conservation with two species in the same cell type.

To better understand how aging impacts retinal function, we concentrated on Müller glia (MG), the principal support cells of the retina, which undergo notable age-related changes in all three species. In aged MG, we observe an evolutionarily-conserved age-dependent increase in activity of gliogenic TFs that are normally active in resting MG – such as Tcf7l1, Sox2, Sox9, Lhx2, Id1, and Nfix – are known or predicted to promote glial quiescence ^9,62–66^(Fig. 4C). This aligns with our previous observation of age-dependent increases in expression of multiple genes specific to resting MG, and suggests that this may reflect homeostatic, anti-inflammatory activity. The age-dependent increase in Srebf1 and Srebf2 activity is consistent with the observed increase in expression of genes associated with lipid metabolism in MG and their known role in age-related diseases such as AMD ^67^. However, Srebf1 also selectively regulates a subset of genes specific to young MG, as do Etv1 and Etv5, which are both FGF-inducible ^68,69^. FoxO family transcription factors, which play an evolutionarily ancient role in regulating cellular metabolism and aging ^70,71^, selectively target not only genes specific to MG but also distinct subsets of genes active in both young and aged rod photoreceptors. Since the anti-inflammatory transcription factor Ppara also shows age-dependent increases in activity, this suggests the existence of a life-long chemoprotective function in rod photoreceptors (Fig. 4D).

This may act to counter the influence of the pro-inflammatory factor Stat1 and the glucocorticoid receptor Nr3c1, which also show increased activity in aging rods. Finally, these data also imply a complex and dynamic age-dependent regulation of rod photoreceptor-specific genes, since Nrl is selectively active in young rods while Crx/otx5a and Neurod1 are more active in aged rods. Since these TFs directly activate rod-specific genes ^72^, this may indicate a progressive age-dependent reorganization of GRNs controlling phototransduction gene expression.

Retinal ganglion and bipolar cells also showed multiple examples of evolutionarily-conserved age-dependent changes in transcription factor activity. In young mouse and human RGCs, we observe increased activity of the homeodomain factor Tfap2d, while in aged RGCs of all three species, we likewise see increased activity of the Kruppel factor Klf7 (Table S9) . Both of these transcription factors are strongly expressed in immature RGCs and are required for RGC development ^73–76^, while Klf7 also promotes RGC survival following optic nerve crush ^77^.

Curiously, Klf7 activity was also upregulated in aged human and mouse bipolar cells. Young bipolar cells of all three species show higher activity of Cux1, which is highly expressed in late-stage RPCs but only weakly expressed in differentiated cells ^62^. Aged mammalian bipolar cells also upregulated activities of several other factors that are predominantly expressed in RPCs, including Tfdp2, Pou2f2, and Zfp148. The significance of this ectopic aging-dependent activity remains unclear. Other changes in transcription factor activity likely reflect broader age-dependent metabolic changes. Like rod photoreceptors, young mammalian RGCs and bipolar cells also upregulated activity of Srebf family factors, while aged mammalian bipolar cells upregulated Hif1a. Young mammalian bipolar cells also showed increased activity of the core circadian regulator Clock (Fig. 4F-H, Table S9).

### Identifying aging-dependent changes in extrinsic signaling in retina and RPE

Since changes in cell-cell signaling are a known hallmark of aging ^1,78,79^, we next used our snRNA-Seq dataset to identify dynamic aging-dependent changes in cell-cell signaling. We used the LRLoop software package, as it produces low false-positive rates ^77^. We first queried all possible signaling interactions between major retinal cell types in zebrafish, mouse, and human in both young and old samples (Fig. 5A). In all cases, we observed more interactions in aged samples, and this was particularly pronounced for interactions between MG and other cell types (Fig. 5B, Table S11), with aged human samples showing by far the largest number of interactions, although this may reflect the overrepresentation of >60 year old human samples (Fig. 5B). Zebrafish showed fewer overall cell-cell interactions than mouse, although MG showed the largest number of interactions, particularly as the sender cell in aged retinas (Fig. 5C). In humans, however, MG were less prominently represented than inner retinal neurons, particularly retinal ganglion cells. Moreover, for all species save humans, rod and cone photoreceptors showed the fewest age-dependent changes in cell-cell communication (Fig. 5C).

**Figure 5:**
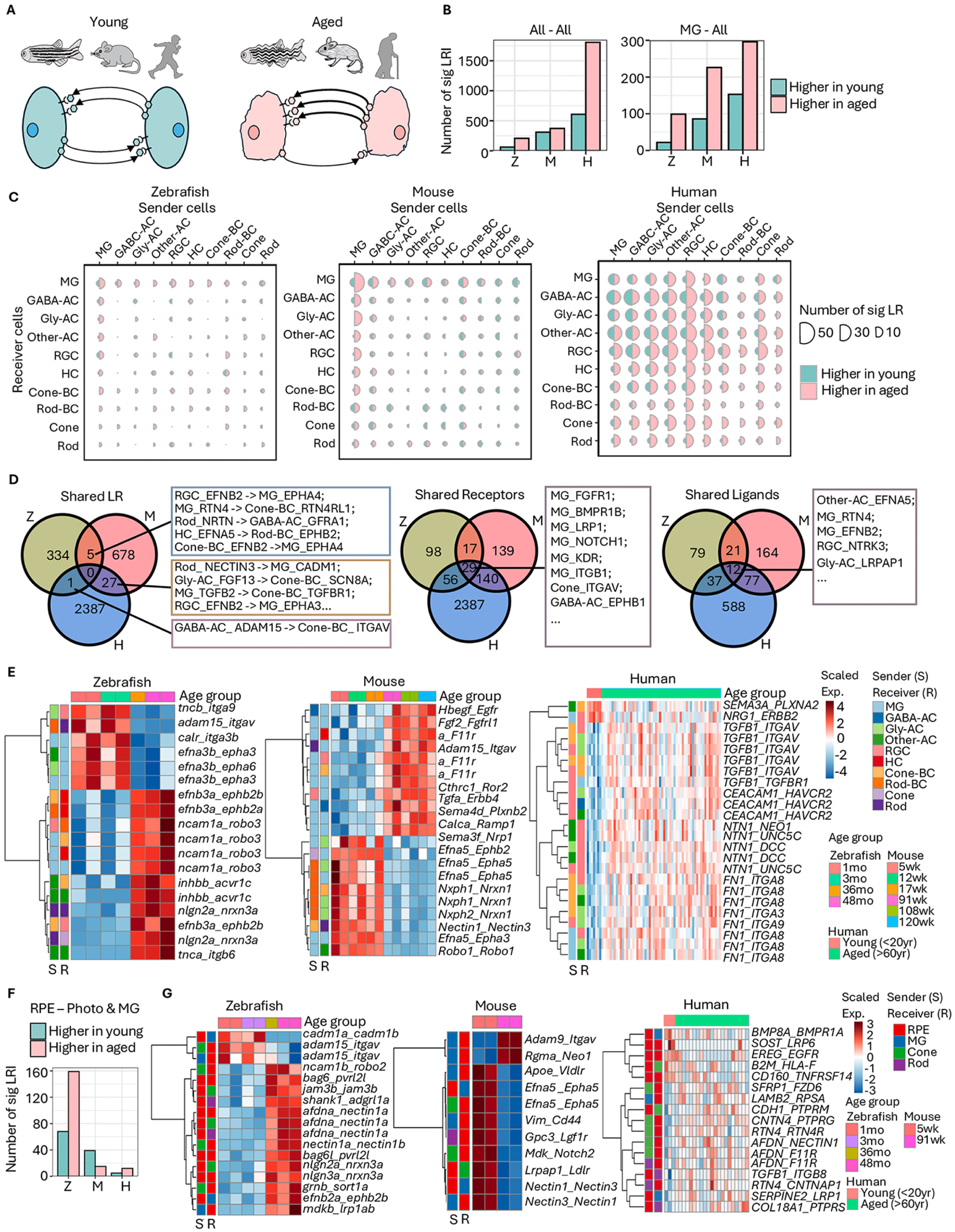
Cell-cell communication network across retinal cell types in zebrafish, mouse and human. (**A**) Schematic overview of the comparison of cell-cell communication networks between young (zebrafish: 1 and 3 months; mouse: 5, 12, and 17 weeks; human: <20 years) and aged (zebrafish: 36 and 48 months; mouse: 91, 108, and 120 weeks; human: >60 years) samples. (**B**) Total number of differentially expressed ligand-receptor interactions (LRI). The cut-off for significantly changed LRI was p-value <0.01. (**C**) Number of differentially expressed LRI between each cell type across the 3 species. (**D**) Shared differentially expressed ligand-receptor (LR) pairs across the 3 species. From left to right: pairs sharing both ligand and receptor genes; pairs sharing only receptor genes; pairs sharing only ligand genes. (**E**) Heatmap of top differentially expressed LR pairs. The left two annotation bars indicate sender (S) and receiver (R) cells, respectively. The interaction scores were scaled across individual samples (row-scaled). (**F**) Total number of differentially expressed LRI between RPE and its interacting cell types. The cut-off for significantly changed LRI was *p* value <0.05. (**G**) Heatmap of top differentially expressed LR pairs for RPE-associated interactions. The left two annotation bars indicate sender (S) and receiver (R) cells respectively. The interaction scores were scaled across individual samples (row-scaled).

The general increase in the complexity of cell-cell signaling in aged retina is consistent with our observation of age-dependent increases in inflammatory signaling, particularly in mice (Fig. 2). To determine whether this accounted for the observed age-dependent increase in cell-cell signaling, we separately analyzed age-dependent changes in cell-cell interactions linked to inflammation (Fig. S8A). All three species showed strong age-dependent increases in inflammatory signaling, with zebrafish and humans also consistently showing age-dependent increases in all other forms of cell-cell signaling, both in all cells and in Muller glia, the retinal cell type that consistently shows the strongest transcriptional response to inflammation (Fig. S8A). Similar patterns of age-dependent cell-cell inflammatory signaling were observed to those of all other signaling modalities. Zebrafish showed the fewest inflammation-associated cell-cell signaling interactions, followed by mice, with humans showing the most. In zebrafish and mice, Muller glia consistently showed the largest number of age-regulated cell-cell signaling interactions, while in humans the most interactions were observed in inner retinal neurons, particularly retinal ganglion cells (Fig. S8B).

When we directly examined the overlap of aging-regulated ligand–receptor pairs across cell types between different species, however, we observed some common pairs, particularly between mouse and human, but did not detect any that were shared by all three species (Fig. 5D). Because related paralogous ligands can often target a given receptor, and vice versa, we separately looked for overlap among receptors and ligands, and identified greater overlap.

Notably, MG in all three species showed age-dependent changes in the signaling activity of FGFR1, BMPR1B, LRP1, and NOTCH1, implying a role in integrating FGF, BMP, Wnt and Notch signaling (Fig. 5D). Ligand-receptor interactions conserved between human and mouse also frequently involved MG as signaling hubs, with rod-derived Nectin3 and RGC-derived Efnb2 respectively targeting glial Cadm1 and Epha3 (Fig. 5D).

The strongest age-dependent interactions were generally species-specific, with MG being heavily represented as signal senders and/or receivers in mouse and zebrafish (Fig. 5E). In young cells, we observed strong and partially-conserved MG-MG signals in both zebrafish (efna3b_epha3) and mice (Efna5_Epha5). MG also consistently showed more injury and inflammation-related receptor ligand pairs (e.g. Fgf2_Fgfrl1 in mice). TGFalpha (Tgfa_Errb4 in mice) and TGFbeta (inhbb_acvr1c in zebrafish and TGFB1_TGFBR1 in humans) both consistently showed higher activity in aged samples. Many ligand-receptor pairs controlling cell adhesion and synaptic maintenance showed dynamic changes in all species, in line with transcriptomic data.

Microglia play important roles in regulating synaptic homeostasis in the brain, and act in conjunction with macrophages to promote inflammatory signaling ^80–84^. Although we profiled relatively few microglia in mice, we were able to capture sufficient numbers in zebrafish and human retina to perform a similar analysis in these species. As in other cell types, more signaling interactions were observed in aged than in young retina in both species, with microglia-Muller glia interactions most prevalent in zebrafish, and microglia-inner retinal neuron interactions more prominent in humans (Fig. S9A,B). Substantial numbers of conserved age-regulated interactions were observed between microglia and retinal neurons, with a notable number involving microglia-cone (NRLGN33-NRXN, MFGE8-ITGAV) or microglia-cone bipolar cells (JAM3-JAM3, NECTIN3-CADM1) signaling (Fig. S9C). The strongest age-regulated interactions in zebrafish included multiple Eph-Ephrin and Slit-Robo interactions between microglia and neurons in aged retina, consistent with potential regulation of synapse number. In humans, in contrast, interactions related to synaptic maintenance, such as Sema-Plexin/Nrp and Ntn-Unc5b were more prominent in young retinas (Fig. S9D).

Finally, given the central role of the RPE in regulating the onset of age-related diseases such as AMD ^85^, we separately analyzed ligand-receptor interactions involving the RPE and adjacent cell types (rods, cones, and MG) by RPE-enriched datasets. In contrast to the whole retina, we observed the largest number of changed interactions in zebrafish and the fewest in humans (Fig. 5F). Few shared interactions were observed (Table S11). As with intraretinal signaling, cell-cell adhesion interactions were prominent in all species, particularly those involving integrins and nectins (Fig. 5G). Aged zebrafish samples showed increased signaling activity by both midkine and *bag6*/*bag6l*, both of which are associated with inflammation and cellular stress. COL18A1_PTPRS are likewise implicated in age-dependent retinal detachment in humans ^86^. Curiously, however, many of these interactions were reduced in aged mice (Vim_Cd44 and Mdk2_Notch2) and humans (B2M_HLA-F and CD160_TNFRSF14).

### Spatial transcriptomic analysis of mouse retinal aging and cell-cell signaling

To validate our snRNA-Seq data and further investigate aging-dependent cell-cell signaling changes spatially, we generated mouse datasets by the Xenium 5k platform to analyze aging-dependent gene expression changes in the mouse retina *in situ*. This retina dataset consists of 4 ages of retina samples, at 5, 12, 55, and 112 weeks, and each age contains two sections (center and peripheral) from one male and one female. Also, each dataset contains 5,000 unique gene probes that extensively overlap with our snRNA-Seq dataset. After clustering and UMAPs analysis, we identified all the major retina cell types and RPE from Xenium datasets (Fig. 6A,B). To compare the snRNA-Seq and Xenium aging datasets, we plotted the gene versus age correlations, and observed positive correlation between aging-dependent changes in gene expression in all major cell types, ranging from 0.22 in Muller glia to 0.05 in RPE (Fig. S10A). High concordance of age-dependent changes was evident upon visual inspection as well as automated signal analysis, particularly for Muller glia (Fig. 6C,D). We next applied the same methodology that was used to generate snRNA-Seq-based cellular aging clocks to the Xenium dataset in order to generate cell-specific aging clocks for each major cell type (Fig. S10B). These showed similar aging-dependent correlations to the snRNA-Seq clocks.

**Figure 6:**
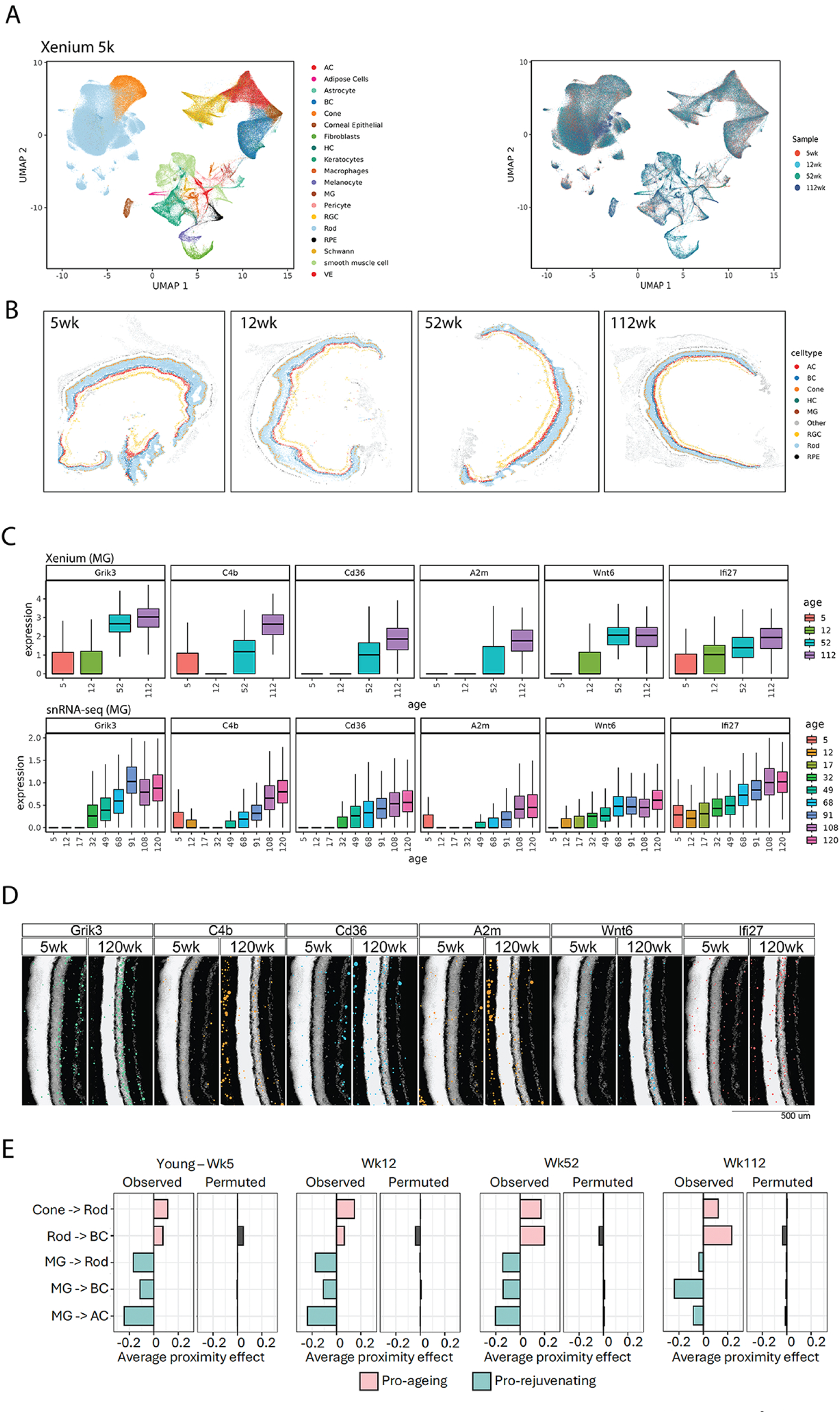
Spatial profiling of aging mouse retina using Xenium5k spatial transcriptomics. (**A**) UMAP embedding of the integrated aging Mouse Xenium dataset, with each spot colored by its assigned cell type (left) and age (right). (**B**) An example of the spatial map of the mouse retina for each age profiled. Cells are colored by eight principal retinal neuron classes—AC (amacrine cells), BC (bipolar cells), Cone, HC (horizontal cells), MG (Müller glia), RGC (retinal ganglion cells), Rod, and RPE (retinal pigment epithelium). (**C**) The boxplots showing examples of consistent aging related genes between Xenium and snRNA-seq datasets in MG cells. The x axis indicates ages of the samples and the y axis the z-score of the gene expression values. (**D**) Xenium images of aging-related gene expression. Scale bar: 500 μm.

To identify robust predictive genes by cellular aging clocks, we examined the overlap of aging-clock features derived from both techniques. To make the aging clocks comparable, we retrained the snRNA-Seq clocks with the same 5000 gene features detected in Xenium datasets. We then compared the gene weights from the clocks by each cell type (Fig. S10C). We identified several strong aging predictor genes with high weights in both models. These genes were also differentially expressed across multiple cell types. Many of the genes with positive weights in multiple cell types were inflammation-associated including *Grik3*, *C4b*, *Ifi27*, *Wnt6*, *A2m*, *Col4a3*, and *Aldoa*. The genes with negative weights were more cell-type-specific, and included *Prdm1* and *Dcx* in rods, *Cxcr4* in MG, *Nefm* in RGCs, and *Pdc* in cones (Fig. S10C).

We next ask whether retinal cells age independently or if specific cell types influence the aging rates of their neighboring cells through intercellular interactions (Fig. 6E). To test this concept, we performed a proximity effect analysis ^87^. First, we applied a regression model to construct aging clocks (i.e., a combination of aging-related genes) for each major retinal cell type, enabling us to predict the “age” of individual cells based on the gene expression profile of this set of genes. Using the Xenium assay, we obtained both gene expression profiles and relative spatial locations of individual cells in the mouse retina. By applying the aging clocks, we predicted the “ages” of individual cells on the Xenium slides. For a given effector cell type, we identified nearby target cells and compared their average predicted ages to those of more distant target cells (Fig. 6E). This approach allowed us to assess the potential influence of proximity on cellular aging within the retina.

With our analysis, we found that cells adjacent to photoreceptors (rods and cones) exhibit accelerated aging compared to distant cells, which suggests that photoreceptors exert a pro-aging effect on neighboring cells. This effect is most pronounced in rod-to-bipolar cell and cone-to-rod cell interactions, and remains consistent across three samples. The effect is statistically significant when compared to permutation controls. Conversely, Muller glia-derived signals rejuvenate neighboring cells, as nearby rods, bipolar, and amacrine cells display younger ages than their counterparts far from Muller glia (Fig. 6E). This analysis suggests that cells are interconnected and that the aging rates are influenced by nearby cells through intercellular interactions.

### Spatially-resolved patterns of aging-dependent changes in cell-cell signaling

While the cell type composition of the retina is relatively uniform in mice over the lifespan, the central and peripheral retina are exposed to substantially different levels of high-energy photon exposure across the lifespan ^88,89^. Furthermore, we already demonstrated that zebrafish exhibit spatially resolved selective aging-dependent expression of *dkk1b* in the peripheral retina (Fig. S2E), while in humans central and peripheral retina show differential vulnerability to age-related diseases such as AMD and glaucoma ^85,90^. Using the Xenium dataset, we systematically investigated differences in age-dependent gene expression and cell-cell signaling between central and peripheral retina. While most aging-regulated genes showed essentially identical patterns of expression across the retina, some of the most strongly age-dependent genes showed dramatic center-periphery differences in expression (Fig. 7A). These include the peripherally-enriched inflammation-associated genes such as *A2m* and *Cd36* in MG, and center-enriched *Grik3* in both MG and bipolar cells (Fig. 7B). *Col4a3* showed a strong age-dependent increase in rods across the retina, but only in peripheral cones. Center-periphery differences were observed for a small number of genes that showed decreased expression in aged retina, most notably including *Efnb2* in MG and amacrine cells (Fig. 7A).

**Figure 7:**
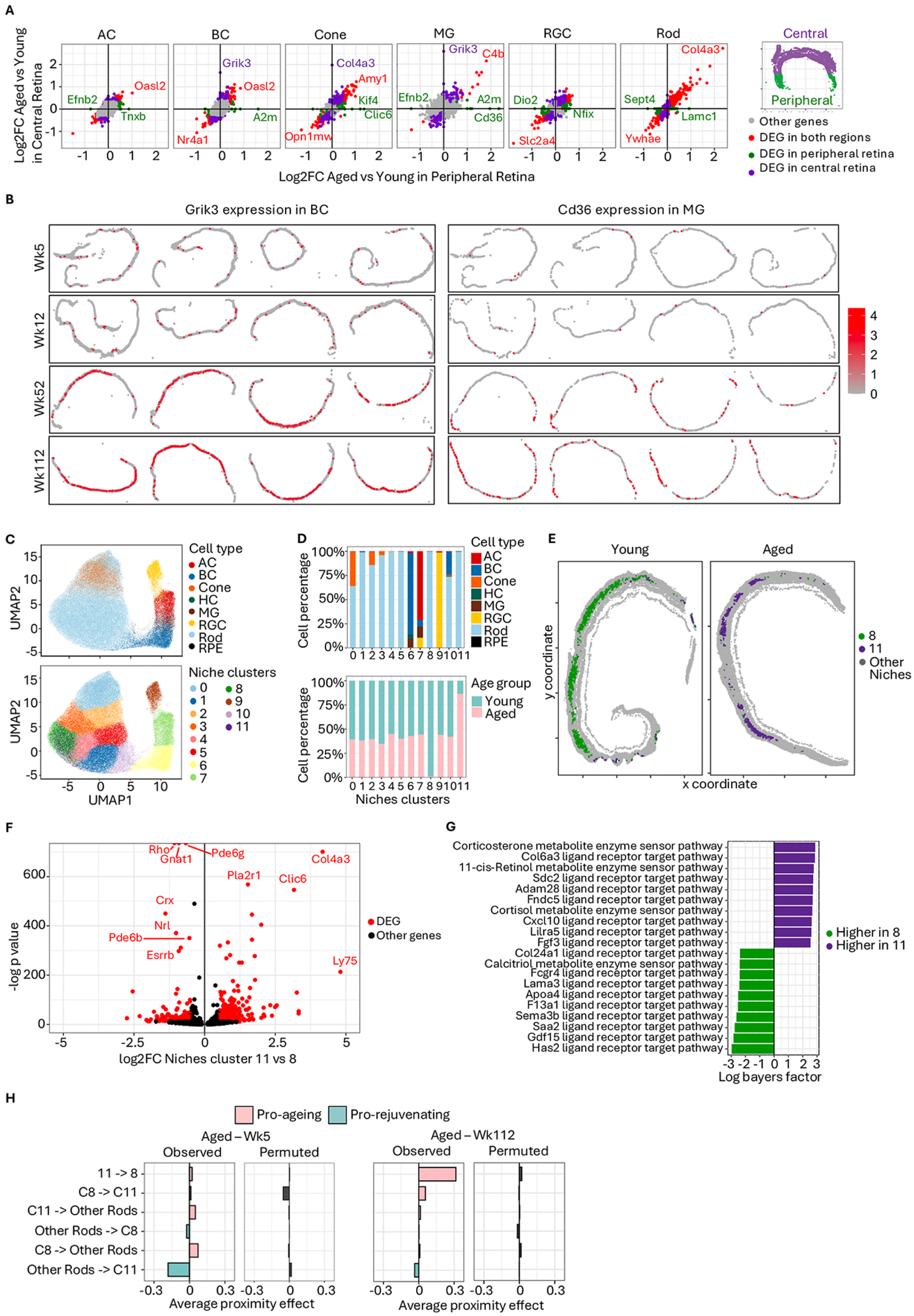
Spatiotemporal cell-cell communication signatures between young and aged mice. (**A**) Log2 fold change of DEGs between aged (112 week) and young (5 week) mice in central and peripheral retina. An example of central and peripheral retina is shown on the right (n=4 for each age group). **(B**) Expression of *Grik3* and *Cd36* gene which showed differential expression patterns between central and peripheral in BC and MG cells during ageing. (**C**) UMAP embeddings of spatial transcriptomes calculated based on cell microenvironment using NicheCompass. Cells are color-coded either by previously annotated cell type (top figure, see Figure 5 for reference) or Niche clusters (bottom figure). (**D**) Top: Proportion of each annotated cell type in each Niche cluster. Bottom: Proportion of aged (112 weeks) and young (5 weeks) cells in each Niche cluster. (**E**) Xenium spatial map of representative young and aged samples. Cells were color-coded by Niche clusters. (**F**) Volcano plot of differential expressed genes (DEG) between aged-specific Niche cluster 11 and young-specific Niche cluster 8. (**G**) Top-enriched cell-cell communication pathways for aged-specific Niche cluster 11 and young-specific Niche cluster 8. (**H**) Cell proximity effect of Niche cluster 11 and 8 on cells in other rod clusters.

To investigate spatial aspects of cell-cell communication, we employed the graph deep-learning framework NicheCompass, which quantitatively characterizes cell clusters (niches) by predicting molecular profiles related to spatially encoded signaling events ^91^. Based on their distinct retinal microenvironment, cells were split into 12 distinct clusters (Figure 7C). Most niches are specific to one cell type, but surprisingly 9 of these niches were essentially specific to rod photoreceptors, which showed the smallest number of aging-dependent cell-cell interactions using LRLoop (Fig. 7D and Fig. 5). Two of these niches – Niche 8 and Niche 11 – were strongly age-dependent, with Niche 8 confined to young cells and Niche 11 overwhelmingly seen in aged cells (Fig. 7D and E). Both of these niches are confined to the central retina (Fig. 7E). Niche 8 is highly enriched for rod-specific phototransduction genes (*Rho*, *Gnat1*, *Pde6g*, *Pde6b*) and transcription factors (*Crx*, *Nrl*, *Esrrb*) relative to Niche 11 (Fig. 7F). Niche 11, in contrast, is enriched for aging-regulated genes such as *Col4a3*, *Ly75*, and *Plr2a1*. Finally, we examined cell-cell signalling pathways predicted to shape the individual niches (Fig. 7G). We observed significantly more cell-cell communication related to corticosterone, FGF and Lilra5 signaling in Niche 11 relative to other Niches, consistent with both increased cellular stress and inflammation associated with cellular damage and aging and, in the case of FGF, homeostatic responses to counteract this ^92–94^. The increased 11-cis retinol-related signaling seen in Niche 11 may reflect compensatory changes in response to reduced expression of phototransduction genes relative to Niche 8. These results imply that retinal cellular aging is not spatially homogenous, and that both pro- and anti-aging extrinsic signals can modulate its rate of progression (Fig. 7H).

### Pro-rejuvenative effect of Muller glia-specific expression of Yamanaka factors in rod photoreceptors and bipolar cells

Low-level or transient expression of the Yamanaka factors Oct4 (Pou5f1), Sox2, and Klf4 has been shown in some contexts to induce cellular rejuvenation without adverse consequences such as dedifferentiation or tumor formation ^95,96^. We directly tested whether constitutive, low level, Muller glia-specific expression of these factors resulted in rejuvenation of any retinal cell types, using the Muller glia-specific inducible Cre line GlastCreER ^65^ in conjunction with Cre-inducible rtTA and doxycycline-inducible *Oct4*, *Sox2*, *Klf4*, and *mCherry*. We induced rtTA expression in *GlastCreER;R26-lsl-rtTA;tetO-Oct4-Sox2-Klf4-mCherry* 12 month old mice using tamoxifen, following which mice were maintained on either control or doxycycline-supplemented chow (Fig. 8A). At one month following treatment, a small number of Sox2-positive Muller glial cell nuclei were displaced into the retinal outer nuclear layer, consistent with limited levels of de-differentiation ^9,97^ (Fig. 8B).

**Figure 8:**
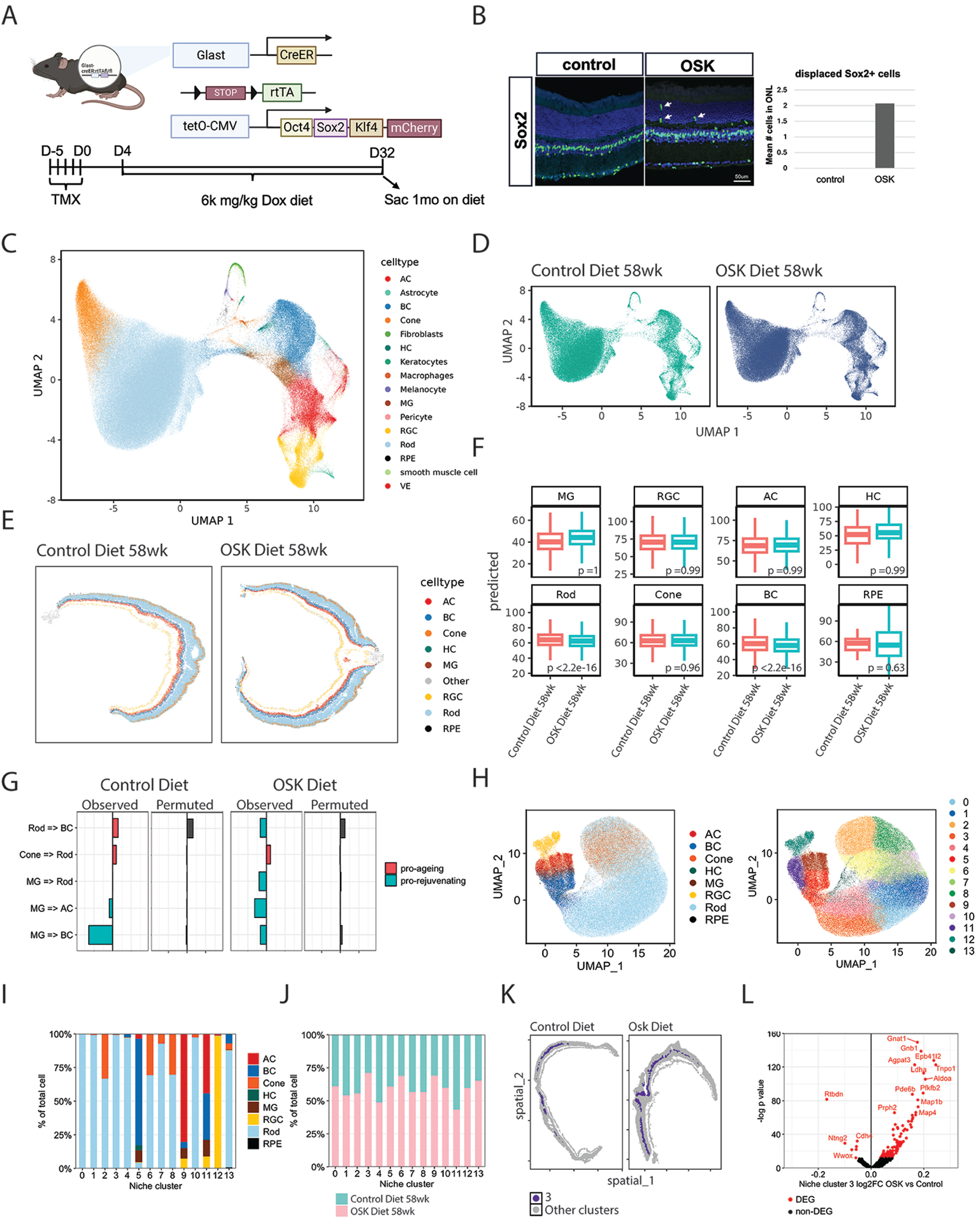
Non-cell autonomous rejuvenative effects induced by Yamanaka factor overexpression in Müller glia. (**A**) Schematic for transgenic Glast:OSK mice and experimental design. (**B**) Left: Immunofluorescence labeling of Sox2 (green). Arrows indicated displaced Müller glia in the outer nuclear layer (ONL). Scale bar represents 50um. Right: Quantification of the mean number of sox2+ displaced Muller glia in the ONL. (**C-D**) UMAP embedding of the integrated aging Mouse OSK overexpression Xenium dataset, with each spot colored by its assigned cell type (**C**) and group (**D**). (**E**) Representative spatial maps of OSK-overexpressing mouse retinas. Cells are colored according to eight principal retinal neuron classes: amacrine cells (AC), bipolar cells (BC), cone photoreceptors (Cone), horizontal cells (HC), Müller glia (MG), retinal ganglion cells (RGC), rod photoreceptors (Rod), and retinal pigment epithelium (RPE). (**F**) Predicted ages were inferred using Xenium-based aging clocks for major retinal neuron populations in control (diet-only) versus OSK-overexpressing samples. The p-value from a one-sided t-test comparing diet-only and diet+OSK groups is shown in each panel. (**G**) Effects of control doxycycline and OSK overexpression on cell-cell signaling. (**H**) UMAP embeddings of Mouse OSK Xenium dataset calculated based on cell microenvironment using NicheCompass. Cells are color-coded either by previously annotated cell type (left figure) or Niche clusters (right figure). (**I**) Proportion of each annotated cell type in each Niche cluster. (**J**) Proportion of cells from Control and OSK mouse for each Niche cluster. (**K**) Xenium spatial map of representative Control and OSK samples. Cells were color-coded by Niche clusters. (**L**) Volcano plot of differential expressed genes (DEG) between Niche cluster 3 cells from OSK mouse and Control mouse.

Doxycycline-treated retinas were analyzed using Xenium 5K spatial transcriptomics (Fig. 8C-E), and cellular age was determined using our Xenium-based aging clock. This revealed that the cellular age of rod photoreceptors and bipolar cells were significantly reduced in OSK-expressing retinas, although that of Muller glia showed no significant decrease. (Fig. 8F).

Doxycycline treatment also enhanced the pro-rejuvenative Muller glia-derived signals targeting rods, amacrine, and bipolar cells (Fig. 8G). Niche analysis of doxycycline-treated samples further identified 13 niche clusters (Fig. 8I). OSK-expressing retinas showed a higher fraction of cells in the rod-specific Niche 3 (Fig. 8J,K). Genes in this cluster were in turn highly enriched for rod-specific genes (Fig. 8L), a pattern which closely resembled genes specific to that rod-specific Niche 8 that is found specifically in young retina (Fig. 7D-G).

## Discussion

Aging-regulated neurodegenerative diseases are a huge public health burden, but the mechanisms by which increased biological age leads to disease development and progression remains unclear. The retina is a particularly tractable system for studying CNS aging, since its anatomy and cell type composition are both relatively simple and evolutionarily well-conserved. Furthermore, age-related neurodegenerative retinal diseases, such as glaucoma and macular degeneration, are highly prevalent in the human population, affecting approximately 1 and 10% of the US population over the age of 50 respectively, with the incidence of both conditions increasing progressively with age ^28,98^. In addition, the retina provides a powerful system to assess the progression of a number of other neurodegenerative diseases, including Alzheimer’s and Parkinson disease ^23,25,26,99,100^.

Here, we use integrated snRNA-Seq and scATAC-Seq analysis to profile aging-regulated changes in gene expression and chromatin accessibility across the natural lifespan in zebrafish, mouse, and human retina, and coupled this with spatial transcriptomic analysis of the naturally aging mouse retina. This has allowed us to identify robust aging-related changes in gene expression of all the major retinal cell types, in all three species. We then used this dataset to generate predictive molecular clocks to measure cellular age, to identify evolutionarily-conserved and species-specific aging-dependent gene regulatory and cell-cell signalling networks, and to identify pro- and anti-aging signaling mechanisms using Xenium-based spatial transcriptomics. Some of the more strongly aging-regulated genes overlap with genes previously identified in smaller scale scRNA-Seq studies of mouse retina – such as *A2m*, *Wnt6* and *Grik3* in Muller glia. These smaller studies also identified a subset of aged rod photoreceptors as expressing lower levels of phototransduction genes, much as we see in Niche 11 in this study ^101,102^. Nonetheless, the large numbers of samples and cells analyzed here, along with the ability to compare these changes across multiple species, make this dataset especially useful in analysis of the landscape of aging-dependent gene regulation.

Correlating our snRNA-Seq data with the aging-regulated genes identified via Xenium spatial transcriptomics, we have identified a core set of aging-regulated cell type and species-specific genes. However, in line with single-cell RNA-Seq studies of other tissues ^4,15,16,48,52,103–106^, we find that the great majority of aging-regulated genes are cell type-specific. Furthermore, direct comparison of aging-dependent changes in identical cell types, revealed that aging-dependent changes are overwhelmingly species-specific, and that the relative fraction of aging-regulated genes differed substantially by cell type across species. This does not result from a broad dysregulation of transcriptional regulation with age, but instead appears to be stereotyped and reproducible. While this seems to be more consistent with a programmed rather than a random mechanism of cellular aging, the lack of evolutionary conservation implies that any such program is idiosyncratic and highly variable across cells and species ^14,16^.

We were nonetheless able to identify a limited number of common pathways and functional categories of aging-regulated genes that were conserved across species. This most notably included increased expression of inflammatory genes in Muller glia and microglia, increased expression of autophagy-associated genes in multiple cell types, and dynamic expression of genes regulating synaptogenesis, cell adhesion, extracellular matrix components, lipid metabolism, and protein degradation. These are all consistent with hallmarks of aging in other tissues. Evolutionarily-conserved age-dependent changes in gene expression that have previously been observed in other tissues include upregulation of class I MHC genes ^107–110^, upregulation of the lipid carrier proteins *Apoe* and *Clu* ^104,111^, and downregulation of genes encoding ribosomal proteins and splicing factors ^112–115^.

Some studies reported a widespread aging-dependent loss of repressive chromatin marks and DNA methylation that leads to a broad increase in chromatin accessibility ^59,116–118^, and we consistently observe an age-dependent increase in chromatin accessibility across all human cell types. However, we do not observe consistent age-dependent changes in chromatin accessibility in any cell type in zebrafish and mice. The reason for this dramatic difference in age-dependent chromatin accessibility is unclear, and requires further investigation.

Reconstructing aging-dependent gene regulatory networks also identified age-dependent changes in the activity of inflammation-associated Stat factors and known aging regulators such as FoxO factors, which broadly promote cellular resilience ^70,71^. However, some of these same factors activated age-dependent genes quite generally – activating different subsets of genes selectively expressed in young and aged cells. Furthermore, we consistently observe an age-dependent increase in the activity of a subset of transcription factors that promote glial quiescence and activate expression of glial-specific genes, including *Nfix, Tcf7l1, Sox9, and Lhx2*. This is also observed in rod photoreceptors and retinal ganglion cells, where age-dependent increases in the activity of transcription factors such as Crx and Klf7 that are prominently expressed in immature postmitotic precursor cells and that drive expression of markers of terminal differentiation of these cell types ^73,119^. This may reflect a homeostatic response to age-dependent increases in inflammatory signaling, which represses expression of phototransduction genes and genes specific to resting glia ^9,120–122^.

This role in maintaining terminal differentiation is consistent with the observation that these same transcription factors also activate selectively genes specific to young cells, and suggests that these may be promising targets for therapies aimed at slowing or reversing the process of cellular aging. We also observe several cases of highly cell- and species-specific patterns of ectopic expression of cell type-specific transcription factors, which are consistent with more restricted dysregulation of gene regulation. These include age-dependent expression of the RGC-specific factor *Pou6f2* in mouse rod and cone photoreceptors, and the glial-specific factor *KLF6* in human rods. This may lead to aberrant expression of target genes that contribute to the process of photoreceptor aging.

While a number of aging-regulator ligands and receptors show conserved activity across all three species, we do not observe any cases where both the ligand and receptor are fully conserved. We find that Muller glia act as core signaling hubs, particularly in zebrafish and mouse, and that cell-cell signaling interactions increase with age across all species and cell types, which partially reflects a global increase in inflammatory signaling. EphrinA-mediated signaling was higher in young samples, as were signals that control synaptic development and maintenance, such as neuroligin, neurexin, and Robo, and semaphorin-mediated signaling. In aged samples, and particularly in human cells, integrin and TGFbeta signaling were prominent. Aging-dependent changes in synaptic organization in the outer plexiform layer was previously noted in mice and observed in this study, and this analysis reveals multiple potential interactions between rods, horizontal cells, and rod bipolar cells that may mediate these age-dependnt changes ^20,44,45^.

Xenium-based spatial transcriptomic analysis not only reveals a common set of cell type-specific genes that are robustly age-dependent across multiple platforms, but unexpectedly identifies spatial heterogeneity in expression patterns of aging-regulated genes. Notably, we observe selective age-dependent upregulation of the inflammatory markers *A2m* and *Cd36* in peripheral retina, while the glutamate receptor *Grik3* is exclusively upregulated in bipolar and MG in the central retina. While *A2m* is known to be upregulated in MG following optic nerve crush ^123^, the significance of these findings is broadly unclear. Since the central retina receives more high-energy photons, we would expect inflammatory signaling to be higher here. Immune cells are known to infiltrate the retina through the ciliary body and iris ^124^, and this raises the question of whether this process might accelerate with age. Previous work demonstrated a role for transient perinatal expression of *Grik3* in mediating light-dependent apoptosis ^125^, but in aged animals *Grik3* is selectively upregulated in the inner retina. Cumulative differences in light-induced glutamatergic signaling over the lifetime may contribute to these regional differences across the mouse retina ^126,127^.

Finally, we applied our molecular aging clock data to the spatial transcriptomic data to both identify pro- and anti-aging patterns of cell-cell signaling among cell types, and to identify the presence of pro- and anti-aging microenvironments among rod photoreceptors. We find that while Muller glia do show the strongest age-dependent increase in inflammatory genes, they also have a strong pro-rejuvenative effect on specific retinal cell types – most notably rods, bipolar, and amacrine cells – which persists throughout life. This is consistent with our prediction that Muller glia serve as signaling hubs to regulate aging-dependent changes in gene expression, particularly in mice, and is in line with recent findings that adult neural stem cells show pro-rejuvenative activity ^87^. We further find that selective, inducible and low level transgenic expression of Yamanaka factors in Muller glia leads to cellular rejuvenation in rod photoreceptors and bipolar cells, as predicted from this analysis. This treatment did not rejuvenate Muller glia themselves, however, and the molecular mechanism mediating this non-cell autonomous rejuvenation remains unclear. These findings suggest that Muller glia, which directly contact every retinal cell type, may eventually prove to be a fruitful target for therapies aimed at rejuvenating the aged retina.

While we observe that rod-bipolar and cone-rod signaling does broadly exert a pro-aging effect, we also identify unexpected heterogeneity among the rods themselves, which are generally considered to be the most molecularly homogeneous retinal cell type. We detect two distinct molecular microenvironments in the retinal outer nuclear layer that are selectively found in young and in aged retinas, respectively. The aged microenvironment, which is associated with a broadly inflammatory profile and with reduced expression of phototransduction genes, spreads progressively inward from the peripheral retina, and promotes conversion of the young microenvironment to an aged state. Yamanaka factor overexpression in Muller glia partially restores the young microenvironment and enhances the expression of phototransduction genes. This not only demonstrates the power of spatial transcriptomic analysis, but identifies new potential therapeutic targets for promoting photoreceptor rejuvenation, thereby maintaining photoreceptor function in age-related photoreceptor dystrophies.

## Acknowledgement

We thank W. Yap for comments on the manuscript. This work was supported by an award from the Milky Way Research Foundation.

## Methods

### Zebrafish maintenance

Zebrafish lines AB and *albino^b^*^4^*^/b^*^4^ ^128^ were raised and maintained at 28.5°C and 30% humidity with a 14h light:10h dark cycle in the Center for Zebrafish Research of the Freimann Life Sciences Center at the University of Notre Dame. All experimental protocols were approved by the University of Notre Dame IACUC committee (21-02-6420 and 21-02-6437) and are compliant with the Association for Research in Vision and Ophthalmology statement for the use of animals in vision research.

### Tissue fixation and cryosectioning – Zebrafish

Tissues for histological analyses were collected at specified time points by euthanizing zebrafish in 1:500 solution of 2-phenoxyethanol followed by ocular enucleation using Dumont#5 and #5/45 forceps (Fine Science Tools). Prior to submersion in fixative, the cornea was poked ventral to the lens with a 30-gauge needle (Fisher Scientific) to promote infiltration of fixative.

Eyes were fixed overnight in 9:1 ethanolic formaldehyde at 4°C and subsequently rehydrated with a series of ethanol dilutions (5 min washes each of 90%, 80%, 70%, 50%, and 30% ethanol in water). Cryoprotection was performed by washing eyes in 1x PBS for 5 min, 5% sucrose (in 1x PBS) 3 times for 5 min each, 30% sucrose (in 1x PBS) overnight at 4°C, and then soaked for 1 to 2 days at 4°C in a mixture of 2 parts Tissue Freezing Medium (TFM; VWR) to 1 part 30% sucrose. Eyes were transferred to 100% TFM and lenses were removed with Dumont #5 forceps after first cutting open the cornea starting at the fixation poke site with Student Vannas Spring scissors (Fine Science Tools). Finally, the eyes were arranged to maintain dorsal-ventral identity in 100% TFM and frozen and stored at -80°C. Frozen tissue blocks were sectioned at -18°C with 14 μm thickness using a Cryostar™ NX50 Cryostat (ThermoScientific). Retinal sections from the dorsal-ventral plane that included the optic nerve head were collected on SuperFrost Plus glass slides (VWR 48311-703), dried for 30 min on at 55°C slide warmer, and stored at -80°C for future immunohistochemical and/or fluorescent *in situ* hybridization staining.

### Immunohistochemistry and *in situ* hybridization staining – Zebrafish

Immunohistochemistry and RNA *in situ* hybridization were performed as previously described ^129,130^. For microglia/macrophage identification, immunolabeling was performed with antibody against Lymphocyte cytosolic protein 1 (Lcp1). Frozen slides were baked at 55°C for 30 min prior to rehydration in 1x PBS for 15 min. Sections were blocked for 2 h in blocking buffer (1x PBS, 10% normal goat serum (SigmaAldrich), 0.2% Triton X-100 (Fisher Scientific), 2% DMSO (Invitrogen)) and then incubated with a rabbit anti-Lcp1 antibody (1:500, GeneTex GTX134697) overnight at room temperature. Slides were washed twice with PBS-Tween20 (0.1%, Fisher Scientific) for 30 min each, then incubated with Alexa Fluor 647-conjugated goat anti-rabbit (1:500, Life Technologies) and DAPI (1:1000, Invitrogen) for 2 h at room temperature followed by two 30 min washes with PBS-Tween20 (0.1%) and, finally, mounted with ProLong Gold Antifade Reagent (Life Technologies) and cover glass (#1.5, VWR).

Single-molecule fluorescence *in situ* hybridization was performed using RNAscope Multiplex Fluorescent v2 Assay (Advanced Cell Diagnostics) according to the protocol for fixed-frozen tissue sample preparation with some modifications and was combined with immunolabeling with antibody against Glial fibrillary acidic protein (Gfap). Frozen sections were washed in PBS for 5 min followed by baking for 1 h at 55°C for 1 h. Tissue sections were post-fixed with 4% paraformaldehyde (Sigma Aldrich) at room temperature for 1 h. Slides were dehydrated for 5 min each in 50%, 70%, and twice in 100% ethanol (Fisher Scientific) and then baked for 1 h at 55°C. Sections were treated with hydrogen peroxide solution (Advanced Cell Diagnostics) for 10 min at room temperature, followed by a distilled water wash prior to boiling in Target Retrieval Buffer (Advanced Cell Diagnostics) for 15 min. The slides were immediately washed with distilled water, dehydrated with 100% ethanol, and dried. Slides were baked for 1 h at 55°C, during which time a hydrophobic barrier was applied to the slide (ImmEdge Hydrophobic Pen; Vector Laboratories), and then dried overnight at room temperature.

Sections were treated with Protease III solution (Advanced Cell Diagnostics) for 30 min at 40°C and washed with distilled water prior to probe incubation at 40°C for 2 h. The following probes (Advanced Cell Diagnostics) were used: Dr-sparc-C1 (cat no. 495811) and Dr-dkk1b-C1 (cat no. 523391). The 3-plex negative control probe mixture was used on a separate slide.

Probe amplification with AMP1, AMP2, and AMP3 proceeded according to manufacturer instructions as did development of the HRP signal with Opal 570 dye (1:1500, Akoya Biosciences). After signal development, slides were washed for 5 min at room temperature in PBS-Tween20 and then processed for immunolabeling as described above. Primary antibody rabbit anti-GFAP (1:500, DAKO Z0334) and secondary antibody AlexaFluor 488-conjugated goat anti-rabbit (1:500, Life Technologies) were used with DAPI nuclear stain.

### Confocal microscopy and image processing – Zebrafish

Images of the GFAP and *sparc* staining were collected as z-stacks of 8 μm thickness with a step size of 1 μm from the dorsal central retina of sections containing optic nerve head using a Nikon A1R inverted confocal microscope equipped with a 40x plan fluor oil immersion objective (numerical aperture 1.3). Images of the Lcp1 staining were collected as z-stacks of 10 μm thickness with a step size of 1 μm from the dorsal central retina of sections containing optic nerve head using a Zeiss LSM 980 inverted confocal microscope equipped with a 40x Plan-Apochromat oil immersion objective (numerical aperture 1.3). Images of the *dkk1b* staining at the ventral periphery were collected as z-stacks of 5 μm thickness with a step size of 0.5 μm and full retinal sections were collected as tiled array (10% overlap) z-stacks of 5 μm thickness with a step size of 1 μm from sections containing optic nerve head using a Zeiss LSM 980 inverted confocal microscope equipped with a 40x Plan-Apochromat oil immersion objective (numerical aperture 1.3).

Identical illumination and camera settings were used to obtain images from each retina within a staining experiment, and identical optimized brightness and contrast were set for all images using ImageJ. Sizing of the full retina tiled arrays was adjusted in Photoshop 2025 (Adobe) to equalize scaling.

### Image quantification and statistical analysis-Zebrafish

Inner plexiform layer (IPL) thickness was measured with the line tool in ImageJ using the GFAP-labeled retina images. An average IPL thickness was calculated from five measurements made along the length of the IPL on each of five images from each age and normalized to the total length of the dorsal retina, which was measured from ciliary marginal zone to optic nerve head on a 10x image of the same retinal section. Lcp-1-positive cells were counted manually with the multipoint tool in ImageJ in all retina layers throughout the 10 μm depth of the z-stack. Counts were normalized to 100 μm retinal length. GraphPad Prism 10 was used to plot the data and perform a one-way ANOVA with Tukey’s post hoc test. The statistical significance is indicated in as * for *p* ≤ 0.05, ** for *p* ≤ 0.01, *** for *p* ≤ 0.001, and **** for *p* < 0.0001.

### Mice

Mice used in these experiments were maintained in accordance with guidelines established by the National Institutes of Health and approved by the Institutional Animal Care and Use Committee (IACUC) at Johns Hopkins School of Medicine (Protocol#MO22022). Mice were housed in a climate controlled facility on a 14/10h light/dark cycle. C57BL/6J mice were used for all experiment and were obtained from Jackson Labs (strain #:000664).

*GlastCreER;Sun1-GFP* mice were obtained from Dr. Jeremy Nathants (cite) and crossed onto *B6;129S4-Gt(ROSA)26Sortm1(rtTA*M2)JaeCol1a1tm7(tetO-Pou5f1,-Klf4,-Sox2,-mCherry)Hoch*/J mice from Jackson Labs (strain #: 034917). These mice were back crossed onto C57BL/6J mice to remove the rtTA gene, and subsequently crossed onto *B6.Cg-Gt(ROSA)26Sor^tm^*^1^*^(rtTA,EGFP)Nagy^/J* from Jackson Labs (strain #: 005670), to generate a tamoxifen and doxycycline inducible line, which we refer to as *GlastCreER;OSK.* Tamoxifen was administered intraperitoneally once per day for 5 consecutive days (1.5mg/dose). Mice were then put on Doxycycline Diet (6kmg/kg) from Teklad Global Rodent Diets (TD.01533) for 4 weeks.

### Sample collection and single-cell multiomics library generation

#### Zebrafish Retina & RPE

Approximately equal numbers of both male and female fish were used at the ages indicated. Zebrafish were euthanized in 1:500 2-phenoxyethanol (Sigma) in fish system water. Zebrafish retinas (8-10 retinas from right eyes of 8-10 fish) were collected at the ages indicated, flash-frozen on dry ice, and stored in a -80°C freezer prior to overnight shipment to Johns Hopkins University for nuclei extraction according to the 10x Multiome ATAC+Gene Expression (GEX) protocol. Briefly, frozen retinal tissues were lysed and nuclei extracted via 10X Genomics Protocol CG000366 Rev C protocol. Nuclei were resuspended in 1 U/µl RNase inhibitor (Promega, N2615) and Diluted Nuclei Buffer (10X Genomics, 2000207). RNA and ATAC libraries were prepared according to the Chromium Next GEM Single Cell Multiome ATAC + Gene Expression protocol (CG0000338 Rev F). Libraries were sequenced via Illumina NovaSeq or NovaseqX Pus targeting 30,000 reads per cell for 10-15000 cells at the Single Cell and transcriptomics sequencing core (STAC) at Johns Hopkins University.

#### Mouse Retina

Retinal tissue from one eye of 2 male and 2 female mice were pooled for each age timepoint and flash frozen on dry ice and stored at -80°C. For all aged mouse multiome libraries, nuclei were extracted per 10X Genomics Protocol CG000366 Rev C protocol from fresh frozen tissue. Nuclei were resuspended in 1 U/µl RNase inhibitor (Promega, N2615) and Diluted Nuclei Buffer (10X Genomics, 2000207). RNA and ATAC libraries were prepared according to the Chromium Next GEM Single Cell Multiome ATAC + Gene Expression protocol (CG0000338 Rev F). Libraries were sequenced via Illumina NovaSeq or NovaseqX Pus targeting 30,000 reads per cell for 10-15000 cells at the Single Cell and transcriptomics sequencing core (STAC) at Johns Hopkins University.

#### Mouse RPE

Aaa For each sample, single-nucleus profiling was performed on RPE isolated from 4 C57BL/6 mice. Two male and 2 female mice were pooled per sample. RPE cells were isolated using a modification of the described protocol ^131^. In brief, mice were sacrificed, and enucleated eyes were placed in Hanks’ Balanced Salt Solution (HBSS) without Ca^2+^ or Mg^2+^ (Gibco, 14175095) on ice. Extraocular tissue was dissected from the orbit, and an incision was made through the cornea to remove the lens. Eyes were individually incubated in 1.5 ml of 1 mg/ml hyaluronidase (Millipore Sigma, H3506) dissolved in HBSS without Ca^2+^ or Mg^2+^ for 45 minutes at 37°C in a 12-well plate. At the end of incubation, samples were transferred to 1.5 ml of HBSS (Gibco, 14025092) and incubated on ice for 30 minutes. The anterior chamber was removed from the eye, and the neuroretina removed without disturbing RPE cells. Next, eye cups were transferred individually to 1.5 ml of 0.05% trypsin with 0.02% EDTA in a 12-well plate, oriented with the anterior side up to facilitate trypsin contact with the RPE cells. Eye cups were incubated at 37°C for 30-60 minutes, until RPE cells began to detach. Eyes were then transferred individually to 1.5 ml of a 20% fetal bovine serum (Gibco, 26140079) solution in HBSS. The eye cups were grasped by the external optic nerve and RPE was dislodged with a gentle shaking motion. Dislodged RPE cells were collected by pipetting and spun down for 10 minutes at 500G at 4°C. RPE cells were then resuspended in 1 ml of cold HBSS and spun down for 5 minutes at 500G at 4°C. The final cell pellet was snap frozen by immersing in an isopentane bath on liquid nitrogen for 1 minute and stored at -80°C until performing single-cell analysis. To isolate nuclei, the described protocol ^132^ was followed with the following modifications. RPE cells were lysed for 10 minutes in a 1X lysis buffer containing a final concentration of 10mM Tris-HCl (Millipore Sigma, T2194), 10 mM NaCl (Millipore Sigma, S6546), 3 mM MgCl_2_ (Millipore Sigma, M1028), 0.1% Tween-20 (Bio-Rad, 1610781), 0.1% Nonidet P40 (Thermo Fisher Scientific, 85124), and 1% bovine serum albumin (Miltenyi Biotec, 130-091-376). Cells were gently triturated every 2 minutes. For ATAC-seq, 0.01% digitonin (Thermo Fisher Scientific, BN2006) was added to the lysis buffer, and 1 mM DTT (Millipore Sigma, 646563) was added to all nuclear isolation solutions. For RNA applications, 1 U/µl RNase inhibitor was added to all nuclear isolation solutions. Three total washes in nuclei wash buffer were performed to promote removal of pigment and cellular debris. Approximately 100000-250000 nuclei were obtained per sample and resuspended in PBS containing 1% bovine serum albumin and 1 U/µl RNase inhibitor (Promega, N2615) for snRNA-seq, or Diluted Nuclei Buffer (10X Genomics, 2000207) for scATAC-seq. snRNA-seq was performed using Chromium Next GEM Single Cell 3’ Reagent Kits v3.1 (10X Genomics) and protocol CG000204 Rev D with dual indexes. snATAC-seq was performed using Chromium Next GEM Single Cell ATAC Reagent Kits v2 (10X Genomics) and protocol CG000496 Rev B. Libraries were sequenced at the Novogene Sequencing Core (Sacramento, CA) on the Illumina NovaSeq X Plus sequencing instrument.

### Sample collection and tissue processing for Xenium

Mouse eyes were enucleated and dissected in ice cold PBS. Eyes were hemisected and lenses removed. Eyes were then embedded in chilled OCT and flash frozen on pre-cooled isopentane and dry ice. Two 14 micron slices were collected from each retina, one peripherally and one centrally, onto a Xenium Prime 5K Mouse Pan Tissue & Pathways Panel slide. Retinas from 1 male and 1 female mouse were collected for each age. Xenium slides were processed at the Single Cell Transcriptomics Core at Johns Hopkins University.

### Mouse tissue processing and Immunohistochemistry

#### Fixation, cryosectioning, and IHC

Mouse eyes were enucleated into ice cold PBS, and cornea and lense were removied. Eyes were fixed in 4% paraformaldehyde (ElectronMicroscopySciences, #15710) for 1h on ice, followed by two 20 minute rinses in 1X PBS. Eyes were then incubated in 30% sucrose over night at 4°C. Eyes were then embedded in OCT and sectioned at 13 micron thickness and dried at 37°C for 15 minutes. Slides were rinsed in 1X PBS and then incubated in primary antibodies diluted in 0.1% TritonX-100 in PBS (PBST) at room temperature for 24h. Primary antibodies: monoclonal rabbit anti-Iba1 (Fujifilm WAKO,012-28521), polyclonal rabbit anti-glial fibrillary acidic protein (GFAP) (Agilent, Z033401-2), monoclonal mouse anti-PKC alpha (Santa Cruz, sc-8393), monoclonal rabbit anti-Cux1+Cux2 (Abcam, ab309139). Slides were washed with 1X PBS to remove excess primary antibodies and were incubated in secondary antibodies in PBST for 10h at room temperature. Slides were mounted with ProLong Gold Antifade Mountant with DAPI (Invitrogen, #P36935) under coverslips (VWR, #,48404-453). Fluorescent Images were captured using Zeiss LSM 700 confocal microscope at the MicFac Microscopy Facility at Johns Hopkins University.

### Quantification & Statistical Analysis

Quantification of immunofluorescence signals was performed using Fiji (ImageJ, NIH). For GFAP, mean intensity density above a set threshold was measured across the entire retinal section. For IBA1, the total number of processes visible in the outer nuclear layer (ONL) were counted. CUX1+2 and PKCα staining was quantified by measuring the mean intensity density above threshold restricted to the ONL. All quantification was performed on comparable regions of interest across samples. Data were compiled and graphed using GraphPad Prism 10.

Statistical analysis was conducted using one-way ANOVA followed by Tukey’s post-hoc test for comparisons across three or more age groups. Results are reported as mean ± standard deviation (SD), with individual data points representing distinct retinas.

### Electroretinography

Full-field flash ERGs were performed as previously described ^133^. Briefly, mice were anesthetized with ketamine (70–100 mg/kg) and xylazine (5–10 mg/kg), and pupils were dilated with a drop of tropicamide and phenylephrine (Alcon, Ft. Worth, TX). Retinal function was assessed using the Celeris ERG system (Diagnosys, Dorset, UK). Light guide electrode-stimulators were placed on each cornea to record bilateral electrical responses, with a platinum ground electrode attached to the tail. Body temperature was maintained at 37°C using the system’s integrated heating pad.

For scotopic ERGs, animals were dark-adapted for at least 12 hours. Light flashes ranging from 0.01 to 1 cd.s/m² were delivered to one eye at a time, using the contralateral eye as reference. The a-wave was defined as the trough of the initial negative deflection, and the b-wave as the amplitude from the a-wave trough to the peak of the subsequent positive deflection. For photopic ERGs, a background light of 30 cd·s/m² was used to suppress rod responses.

Cone-mediated responses were elicited using flash intensities of 3–10 cd.s/m² above background, and flicker responses were recorded at 10 and 30 Hz.

### Xenium Segmentation

To improve the accuracy of cell segmentation in Xenium spatial transcriptomics data, we re-segmented the retinal tissue using high-resolution fluorescent imaging. Tissue sections were imaged with the DAPI channel on a ZEISS Apotome 3 fluorescence microscope (Carl Zeiss Microscopy) using a 20X objective. Z-stack images were acquired across the full thickness of the section with a z-step size of 2.0 µm. A 2D maximum intensity projection was generated from each z-stack using ZEISS Zen software (Carl Zeiss Microscopy), producing high-resolution DAPI images that captured nuclear architecture throughout the tissue.

The 2D projection DAPI images were aligned to the original Xenium DAPI images using the image registration function in Xenium Explorer 2.0 (10x Genomics), generating alignment matrices. Alignment accuracy was verified visually by comparing prominent anatomical landmarks and nuclear contours. Custom Python code was used to apply these alignment matrices to transform the projection images (10.5281/zenodo.15777981).

The aligned images were imported into QuPath 0.5.0 for layer-specific segmentation. Using the cell detection classifier tool, we trained custom classifiers to segment nuclei within the outer nuclear layer (ONL), inner nuclear layer (INL), and ganglion cell layer (GCL) ^134^. Due to the distinct compactness of nuclei in the ONL, segmentation for this layer was performed separately using manual selection to ensure accurate annotation.

Nuclei were detected using the Positive Cell Detection approach with the following parameters:

- ONL: Background Radius: 1 µm, Sigma: 1 µm, Minimum Nuclear Area: 4 µm², Maximum Nuclear Area: 50 µm², Threshold: 600, Cell Expansion: 3 µm, Pixel Size: 0.2125 µm
- INL + GCL: Background Radius: 4 µm, Sigma: 1.8 µm, Minimum Nuclear Area: 10 µm², Maximum Nuclear Area: 60 µm², Threshold: 400, Cell Expansion: 3 µm, Pixel Size: 0.2125 µm Transcripts were re-assigned to cells using the custom segmentation masks via the import-segmentation pipeline in Xenium Ranger 2.0 (10x Genomics), generating an updated cell-by-gene count matrix for downstream analysis. For cells located in extra-retinal regions (e.g., retinal pigment epithelium, choroid), the original Xenium segmentation output was retained for transcript-to-cell assignment.

### Single-cell multi-omics preprocessing

Raw multi-omic sequencing files for zebrafish and mouse were demultiplexed and processed with Cell Ranger ARC, aligning RNA reads to Danio rerio (DanRer11) and Mus musculus (mm10) reference genomes to generate cell-by-gene UMI count matrices. Human aging single-cell datasets were retrieved from the CellXGene portal, and their raw count matrices were extracted by Scanpy.

### Filtering barcode doublets and low-quality cells

For scRNA-seq, low-quality cells were excluded by filtering out barcodes with fewer than 500 or more than 50,000 RNA UMIs. To remove doublets, we applied an iterative Scrublet workflow^135^ per sample: each cell’s doublet score was computed, cells scoring > 0.5 were removed, and Scrublet was rerun on the remaining profiles until the estimated doublet rate fell below 5 %. To characterize ambient RNA contamination, droplets with fewer than 20 total UMIs were classified as “empty droplets”. UMI counts from all empty droplets were summed per gene, and each gene’s ambient fraction was calculated as its share of the total ambient UMIs. Genes with an ambient fraction > 0.5 % were flagged as ambient contamination markers. These genes and mitochondrial genes are removed in the downstream analysis.

For snATAC-seq, fragment files were imported into ArchR ^135,136^ using default settings, retaining only those barcodes that passed scRNA-seq quality control. We further excluded any cells exhibiting a doublet enrichment score > 2. All filtered scRNA-seq and snATAC-seq datasets were carried forward for downstream analyses.

### Clustering, visualization, and identification of cell types

Raw count matrices from all aging snRNA-seq samples were imported into Scanpy (v1.9)^137^ and concatenated into a single AnnData object, with each cell’s sample recorded in the batch field. Counts were normalized to 10,000 per cell (sc.pp.normalize_total), log-transformed (sc.pp.log1p), and the top 2,000 highly variable genes were identified across batches (sc.pp.highly_variable_genes, flavor = “cell_ranger”). These genes were scaled to zero mean and unit variance (sc.pp.scale), and dimensionality was reduced by PCA on the HVG space (sc.tl.pca, n_comps = 50). To integrate these samples, Harmony (via harmonypy v0.0.14) ^138^was applied to the PCA embeddings with batch as the covariate, and the adjusted PCs were retained. A neighborhood graph was built on the first 30 Harmony-corrected components (sc.pp.neighbors, n_pcs = 30), and two-dimensional UMAP coordinates were computed (sc.tl.umap, min_dist = 0.5, spread = 1.0) for visualization across aging timepoints. The integrated snRNA-seq data were annotated using CellAnn^139^. Briefly, we use the average expression values for each cluster as input for CellAnn, then we select the retina datasets from the database and annotate the cell type for each cluster by combining CellAnn results.

For snATAC-seq of Zebrafish, Mouse, and Human, a unified tile matrix and gene-score matrix were computed across all samples using ArchR. Dimensionality reduction and batch correction were performed with ArchR’s iterative LSI and Harmony workflow: initially, addIterativeLSI was applied to the tile matrix (varFeatures = 50,000; iterations = 2) to capture major sources of variation. Subsequently, addHarmony was executed on the LSI embeddings, incorporating the sample field (representing age) as a covariate. Harmony-corrected dimensions were utilized to construct a shared nearest-neighbor graph (addClusters, resolution = 0.8) and generate two-dimensional UMAP embeddings (addUMAP, reduction = "Harmony", nNeighbors = 30, minDist = 0.5). Cell type annotations for Zebrafish and Mouse in the snATAC-seq data were transferred from their corresponding cells from snRNA-seq. For Human datasets, integration of scRNA-seq and scATAC-seq data from the same donor was initially performed using ArchR’s IntegrateData function. Subsequently, cell type annotations derived from Human scRNA-seq were transferred to the corresponding scATAC-seq data using ArchR’s TransferData function.

### Generating fixed-width and non-overlapping peaks

For Zebrafish, Mouse, and Human scATAC-seq samples, fixed-width 501 bp peaks were called for each cell type using MACS2 within the ArchR package. Subsequently, an iterative merging approach was employed to consolidate overlapping peak sets while preserving the most significant peaks. This merging process utilized the "addReproduciblePeakSet" function in ArchR. Following peak merging, peak matrices were constructed for each dataset using the "addPeakMatrix" function with parameters set to ceiling = 4 and binarize = FALSE.

### Identification of aging-related genes

To identify aging-related differential genes, we first calculated pseudo bulk gene expression profiles for each cell type at various chronological age points. For Zebrafish and Mouse datasets, pseudo bulk gene expressions were aggregated, normalized to a scale factor of 1e4, and log2-transformed. Gene expressions were then standardized to z-scores based on chronological age. For human datasets, pseudo bulk gene expression profiles were computed for each donor. Similarly processed, raw counts were aggregated and normalized to 1e4. Due to donor variability, we defined 8 age intervals and calculated gene expressions as the average across all donors within each interval. These profiles were log2-transformed and standardized to z-scores across all intervals.

Following pseudo bulk computation and z-score normalization, we assessed each gene’s expression pattern across age intervals using generalized additive models (GAM) in R. The GAM method determined significant variation in gene expression with chronological age (p-value < 0.05), identifying aging-related differential genes. Subsequently, we explored the expression patterns of these genes using k-means clustering across age intervals. This approach grouped genes with similar age-related expression trends and visualized results using heatmaps. For Zebrafish and Mouse datasets, k-means clustering employed 8 clusters, while for Human datasets, 16 clusters accommodated donor heterogeneity. After clustering, we manually refined results by excluding 4 clusters with discontinuous age-related expression trends in Human datasets. Finally, based on k-means clustering and heatmap visualization, we categorized clusters into "aged," "young," and "middle-aged" based on distinct patterns of age-related gene expression changes.

### Building cell-type-specific aging models

The aging clock was developed by beginning with the application of k-nearest neighbors (KNN) smoothing to single-cell gene expression profiles. Different k-values were used—k=3 for microglia and RPE due to their lower cell counts, and k=15 for other cell types—to enhance signal clarity and reduce noise. This smoothing process was conducted separately for each sample to ensure optimal data preprocessing.

Following this, pseudo bulk gene expression data across different aging time points was analyzed. The objective was to determine correlations between gene expression levels and chronological age. Genes showing correlations greater than 0.3 or less than -0.3 were identified as age-related genes. Known aging-related genes from databases like Aging Map <u>(Mao et al. 2023)</u> and SENCID <u>(Tao et al. 2024)</u> were also integrated to enrich the feature set.

Subsequently, the dataset was partitioned into 30% test and 70% training sets using stratified sampling. For Zebrafish and Mouse datasets, stratification was based on time points, while for Human data, it was based on donors. To manage dataset size effectively, the maximum number of cells per cell type per donor was capped at 750 during sampling.

Elastic Net regression model parameters were optimized and the final aging clock model was trained using the “glmnet" package in R. The “glmnet” function combines both Lasso (L1) and Ridge (L2) regularization to enhance model robustness and prevent overfitting in high-dimensional datasets, such as gene expression profiles across aging time points. During training, we utilized the cv.glmnet function for cross-validation to identify the optimal regularization parameters (alpha and lambda) that minimize prediction error while maintaining model interpretability. Additionally, each cell’s contribution to the model was weighted based on its respective age interval, ensuring that the model captured age-related gene expression changes accurately. Evaluation of the aging clock model’s performance was conducted using the “predict” function on the remaining 30% test data. This step assessed the model’s ability to generalize across different species-specific datasets, validating its predictive accuracy and reliability in inferring biological age from gene expression profiles.

To extend the application of the aging clock for predicting cellular ages in the Xenium dataset, the approach was further refined. Initially, a common set of genes between the training scRNA-seq atlas and the Xenium panel was identified by intersecting feature lists. Using this overlapping gene set, the aging clock models were retrained, following the same log-normalization and scaling procedures as in the original workflow. Hyperparameters for elastic-net regression were fine-tuned via five-fold cross-validation on the scRNA-seq training cohort. Finally, these refined models were applied to each spatially resolved "cell" (spot) in the Xenium data to infer predicted biological ages. Evaluation included examining age distributions across experimental groups and comparing predictions with those from the full-gene-set aging clock.

### Inferring aging-related GRNs from multi-omics

Gene regulatory networks (GRNs) were constructed independently for zebrafish, mouse, and human using the same method as before with minor modifications.^58,138^ Then the aging-related GRNs for each species were subtracted from the total GRNs if their regulated target genes are aging clock genes identified in figure3.

1. Identifying cis-regulatory elements:

Peaks were categorized into three types around target genes: TSS Peaks within a 1 kb region surrounding the transcription start site (TSS), Gene Body Peaks within the gene body with PtoG correlation > 0.25 and FDR < 0.05, and Intergenic Peaks within 500 kb of TSS but not overlapping with other genes’ TSS or gene body regions, also with PtoG correlation > 0.25 and FDR < 0.05.

2. Predicting TF binding sites:

Cell type-specific TF binding sites were predicted based on TF expression levels, motif matching (TRANSFAC2018 and Cis-BP), and footprint scores. TFs with low average expression (< 0.1) were excluded. Binding sites for expressed TFs were identified using motifmatchr::matchMotifs function (p-value threshold = 5e-5). Footprint scores, derived from merged scATAC-Seq signals using TOBIAS software, assessed Tn5 insertion sites normalized across center and flanking regions.

3. TF-target Correlation:

TF-target regulatory scores were computed using the GBM-based GRN inference algorithm from the arboreto package (grnboost2 function) with all the single cells expression matrix in each species. Pearson correlations were also calculated for each TF-target pair. Interactions were categorized as positive (importance score > 0.1 and Pearson correlation > 0.05) or negative (importance score > 0.1 and Pearson correlation < -0.05).

4. Construction of GRNs:

Cell type-specific regulons were constructed by integrating results from Steps 1–3. TF–peak– target regulons were classified as activating or repressive based on TF–target correlations. Duplicated regulons were removed before downstream analyses. TF–target regulons were further analyzed to identify transcription factors promoting cone cell development.

5. Identification of aging DEGs associated regulons:

The aging-associated regulons are classified into four distinct categories based on the changing tendencies of target genes with age and the type of TF-gene interactions: Activate old regulons where TFs activate genes, leading to increased expression over time; Activate young regulons where TFs activate genes, resulting in decreased expression with age; Repress old regulons where TFs repress genes, causing increased expression with age; and Repress young regulons where TFs repress genes, leading to decreased expression over time.

6. Identification of key TFs in 4 classes of aging GRNs.

To identify key activators (TFs) in the aging GRNs, we calculate the coverage score and enrichment p-value for each TF. The coverage score is computed as Noverlap / NTFtotal, where Noverlap represents the number of target genes shared between the TF and aging GRNs, and NTFtotal denotes the total number of target genes for the TF across all GRNs. To assess whether a given TF is specifically enriched in aging GRNs, we employ a hypergeometric test using the "phyper" function in R. Here, the "population" consists of all TF target genes across the total GRNs, the "sample" includes TF target genes from aging GRNs, and "successes" are defined as genes present in both aging GRNs and the TF’s target genes from total GRNs. Finally, TFs with a p-value < 0.05 and coverage > 0.01 are identified as key TFs influencing the aging gene regulatory networks.

### Xenium spatial transcriptomics analysis

Xenium 5k spatial transcriptomics data were processed in Seurat v5 ^58,138,140^ using a standardized pipeline. For each sample, the Cell Ranger spatial outputs were loaded into a Seurat object via the LoadXenium function. Cells with fewer than 100 total counts were excluded during quality control. Expression values were then normalized and variance-stabilized with SCTransform (default settings). The four samples were integrated using IntegrateLayers with the RPCAIntegration method on the SCT-normalized matrices. A shared nearest-neighbor graph was built from the first 30 principal components, and clusters were identified by the Louvain algorithm (FindClusters, resolution = 2). Two-dimensional UMAP embeddings were computed using PCs 1–30. Finally, each cluster was manually annotated by inspecting the expression of well-characterized retinal marker genes.

### GO and KEGG analysis

Significantly enriched Gene Ontology (GO) Biological Process and KEGG pathway terms were identified among: differentially expressed aging genes (young vs. old) and aging-clock feature genes (positive vs. negative weights). Enrichment analysis was conducted using the enrichGO and enrichKEGG functions from the clusterProfiler R package ^58,138,140,141^, with a significance cutoff of p < 0.05. To reveal evolutionarily conserved processes, enriched GO and KEGG terms from zebrafish, mouse, and human were aligned across species manually.

### Building cell-cell communication networks

The cell-cell communication analysis was done using the LRLoop method on snRNA-seq datasets derived from zebrafish, mouse, and human retinas to investigate age-related changes for intercellular interactions among retinal cell types ^142^. In brief, samples from each species were divided by cell type, and LRLoop was applied on every pairwise combination of cell types to identify enriched ligand-receptor pairs. Each individual sample was specified as the condition of interest using the ’condition’ parameter in LRLoop, enabling the calculation of activity score for each ligand-receptor genes (LR) pair within each individual sample. Samples were then categorized into young (zebrafish: 1 and 3 months; mouse: 5, 12, and 17 weeks; human: <20 years) and aged (zebrafish: 36 and 48 months; mouse: 91, 108, and 120 weeks; human: >80 years). LR activity between age groups were statistically compared using ANOVA, adjusting for biological replicates in zebrafish and mice, and sex differences in humans. A *p* value of 0.01 was set as cut-off for differential expressed LR (DELR). Analysis of cell-cell communication involving RPE cells was conducted separately due to the limited number of samples containing sufficient RPE cells. A *p* value of 0.05 was set as a cut-off for DELR for RPE-related cell-cell communication.

### Cell proximity effect

The cell proximity effect was estimated using a previously published python package ^87^. In brief, we utilized our clock-estimated cellular ages to model interactions between neighboring cell types, quantifying their pro-aging or rejuvenating effects. Specifically, we assessed the influence of each effector cell type on the transcriptomic aging of a target cell type by comparing target cells located near effector cells with those positioned further away. The Euclidean distance between cells was computed between the centroid of each cell. We computed distance cutoffs for each effector-target cell type by calling nearby cells as the average of the median neighbour-neighbour distances for each sample (Yang_SupplementaryFiles_Cellproximity_distance_Cut-off). For each cell, we calculated the shortest Euclidean distance to effector cells, labeling target cells below the cutoff as ’near’ and those beyond as ’far’. Matched sets of ’near’ and ’far’ cells were combined to calculate proximity effects using Cohen’s d, comparing age acceleration between ’near’ and ’far’ cells. Comparisons with fewer than 50 cells per group were excluded. Positive values indicated pro-aging effects, negative values indicated rejuvenating effects. Normalized proximity frequency and statistical significance (Student’s t-test) were also computed. A *p* value of 0.05 was set as cut-off for significant pro-aging or rejuvenating effects.

### Spatiotemporal aging signatures

Each retina Xenium slice was split into two regions, central and peripheral retina, to identify region-specific aging signature genes. Differential expression analysis was applied between aged (112 week) and young (5 week) mice on central and peripheral retina respectively. An FDR-adjusted *p* value of 0.05 was set as cut-off for differential expressed genes (DEG).

To further investigate spatiotemporal cell-cell communication signatures, we applied NicheCompass ^91^ to our spatial transcriptomic dataset from young (5 week) and aged (112 week) mice. NicheCompass is a recently developed deep-learning method designed for modeling intercellular interactions by explicitly predicting molecular profiles of cells and their neighbors in relation to specific signaling events, allowing niche identification and characterization based on scoring of pathway usage in cellular microenvironments ^91^. In brief, we applied NicheCompass on our spatial dataset of (2 age group x 4 replicates) with individual sample set as covariant. Model was trained with following hyperparameters: n_neighbour = 4, edge_batch_size = 4096, conv_layer_encoder = "gcnconv", active_gp_thresh_ratio = 0.01, n_epochs = 400, learning_rate = 0.0005. The training was performed on a NVIDIA-Tesla-T4 16GB GPU. Learned embedding of each cell and gene expression matrix were transferred into R package Seurat v5 (ref) for downstream analysis.

## Data availability

Raw and processed single-cell multi-omics and spatial transcriptomics data have been deposited in the Gene Expression Omnibus under accession number GSE307031. Custom analysis code used in this study is available at https://github.com/lp871/Aging_Project_2025

## Supplemental Figures

**Supplemental Figure 1: SnATAC-Seq analysis of naturally aging retina in zebrafish, mouse, and human.**

**(A)** UMAP embeddings of integrated single-cell ATAC-seq for each species. Cells are colored by their major annotated classes: Müller glia (MG), amacrine cells (AC), cone photoreceptors (Cones), retinal ganglion cells (RGC), horizontal cells (HC), rod photoreceptors (Rods), bipolar cells (BC), retinal pigment epithelium (RPE), microglia, and (in human only) astrocytes. **(B)** Cell-type composition across ages. Stacked bar charts display the absolute number of cells in each major class at each time point for zebrafish, mouse, and human. The x-axis shows chronological age; the y-axis shows total cell counts per class. **(C)** Conserved, cell-type-specific accessible genes across species. Dot plots report, for zebrafish, mouse, and human, the average accessibility (“activity”) of each marker gene (color scale) and the fraction of cells in that type with accessible chromatin at that locus (dot size). Selected transcription factors and canonical markers are listed on the y-axis. **(D)** Dendrogram of ATAC-homology among retinal cell types. Hierarchical clustering based on genome-wide chromatin accessibility distances places homologous cell types closer together. Branch lengths indicate global distance; inner rings are colored by species. Cell-type labels surround the outer ring.

**Supplemental Figure 2: Histological markers of aging zebrafish retina.**

**(A)** MG labeling in aging retinas immunostained for GFAP (green) and counterstained with DAPI (blue). Scale bar is 25 μm. ONL, outer nuclear layer; INL, inner nuclear layer; GCL, ganglion cell layer. (**B**) Quantification of inner plexiform layer (IPL) thickness by age in the central dorsal retina of sections containing the optic nerve head. *n*=5 retinas. Data presented as mean ± SD. Statistical comparison by one-way ANOVA with Tukey’s post hoc test. *, *p*≤0.05, **, *p*≤0.01; ****, *p*<0.0001. (**C**) Microglia labeling in aging retinas immunostained for Lcp1 (grayscale) and counterstained with DAPI (blue). Scale bar is 25 μm.(**D**) MG expression from snRNA-seq data of *sparc* and *dkk1b* in aging zebrafish retinas. (**E, E’**) RNA *in situ* hybridization with probe against zebrafish *sparc* (grayscale) in aging zebrafish retinas (**E)** counterstained with DAPI (blue) or (**E’**) immunolabeled for GFAP (green). Scale bar is 25 μm. (**F, F’, F’’**) RNA *in situ* hybridization with probe against zebrafish *dkk1b* (grayscale) in aging zebrafish retinas (**F, F’**) counterstained with DAPI (blue) or (**F’’**) immunolabeled for GFAP (green). Tiled arrays in (**F**) show full retinal sections along the dorsal-ventral plane. Scale bar in (**F)** is 250 μm and in **(F’**) in 25 μm.

**Supplemental Figure 3: Immunohistochemical assessment of mouse retinal aging.** (**A**) Immunofluorescence labeling of GFAP (green) and DAPI (blue). (**B**) Quantification of mean fluorescence intensity density ± SD of GFAP expression throughout all retinal layers. (**C**) Immunofluorescence labeling of Iba1 (green) and DAPI (blue). White arrowheads indicate microglial processes in the ONL. (**D**) Quantification of total number of microglial processes in the ONL. (**E**) Immunofluorescence labeling of Cux1+Cux2 (green) and DAPI (blue). (**F**) Quantification of mean fluorescence intensity density ± SD of Cux1+Cux2 expression restricted to the ONL. (**G**) Immunofluorescence labeling of PKC-alpha (red) and DAPI (blue). (**H**) Quantification of mean fluorescence intensity density ± SD of PKC-alpha expression restricted to the ONL. (**I**) Quantification of scotopic a-wave ERG. (**J**) Quantification of scotopic b-wave ERG. (**K**) Quantification of photopic b-wave ERG. (**L**) Quantification of photopic b-wave ERG flicker at 10Hz and 30 Hz. Scale bar = 50μm. Significance was determined via one-way ANOVA with Tukey’s multiple comparison test: *p < 0.05. Each data point was calculated from an individual retina. ONL, outer nuclear layer; INL, inner nuclear layer; GCL, ganglion cell layer.

**Supplemental Figure 4: GO and KEGG terms enriched in aging-related genes in zebrafish, mouse and human.** (**A**) Dot-plot of enriched GO/KEGG pathways in young (left) vs. old (right) zebrafish retinal cells. Dot size reflects the count of aging-related genes in each term; color indicates enrichment strength (–log_10_P-value). (**B**) Dot-plot of enriched GO/KEGG pathways in young (left) vs. old (right) mouse retinal cells. Dot size corresponds to the number of aging-associated genes per term; color denotes the enrichment score (–log_10_P-value). (**C**) Dot-plot of enriched GO/KEGG pathways in young (left) vs. old (right) human retinal cells. Dot diameter represents the number of aging-related genes in each term; color scale shows enrichment significance (–log_10_P-value).

**Supplemental Figure 5: Species and cell type-dependent enrichment of senescence-associated genes.** Dot plot of enriched senescence-associated gene terms in aged zebrafish, mouse, and human retinal cells. Dot size reflects the number of aging-related senescence genes in each term, and color indicates the enrichment strength (–log_10_P-value).

**Supplemental Figure 6: Sex-dependent differences in expression of aging-regulated genes in human retina.** (**A**) Heatmaps of shared aging-regulated genes in human samples (male vs. female). Each row corresponds to an aging-associated gene; each column represents a defined age interval. Cell colors indicate the Z-score–normalized expression level of each gene across those age intervals. (**B**) Heatmaps of sex-divergent aging-regulated genes in human samples. As above, rows are aging-associated genes that differ between males and females; columns are age intervals. Colors denote the gene’s Z-score across ages. (**C**) Bar plots of the top enriched GO terms among shared aging-regulated genes (male vs. female). Top panel shows terms enriched in younger samples of both sexes; bottom panel shows terms enriched in older samples. The x-axis shows -log_10_(P-value) for each GO term. (D) Bar plots of the top enriched GO terms are shown for sex-divergent aging-regulated genes. Top panel bars correspond to terms enriched in young females and old males; bottom panel corresponds to terms enriched in old females and young males. The x-axis shows -log_10_(P-value) for each GO term.

**Supplemental Figure 7: Age-dependent changes in differentially accessible chromatin regions (DAR) by cell type and species.** Volcano plots showing the DARs for each cell type in zebrafish, mouse, and human between old and young groups. Each point represents an accessible peak, with the x-axis showing log₂(fold change) (old / young) and the y-axis showing –log10(p-value). The total number of significantly differential peaks (FDR < 0.05) for each plot is indicated above the panel.

**Supplemental Figure 8: Inflammation-associated aging-regulated changes in cell-cell signaling.** (**A**) Total number of Inflammation-associated (Inflam) ligand-regulated interactions which significantly changed between young and aged individuals. (**B**) Number of Inflammation-associated (Inflam) ligand-regulated interactions which significantly changed between young and aged individuals. Numbers were shown for every cell type combination separately.

**Supplemental Figure 9: Aging-dependent changes in cell-cell signaling in microglia.** (**A**) Total number of significantly changed ligand-receptor interactions between young and old individuals for zebrafish and humans. (**B**) Number of significantly changed ligand-receptor interactions between young and old individuals grouped by different cell type combinations. (**C**) Shared ligand-receptor interactions, shared ligands, or shared receptors between zebrafish and human. (**D**) Heatmap of top differentially regulated ligand-receptor interactions between microglia and other cell types.

**Supplemental Figure 10: Comparison of aging-related DEGs and aging clocks between Xenium and snRNA-seq.** (**A**) Scatter plots display the correlation coefficients between gene expression and chronological age for all 5,000 genes in two independent datasets: Xenium (y-axis) versus snRNA-seq (x-axis). Each point represents one gene. The Pearson correlation coefficient between the 2 correlation coefficients is indicated above each panel, summarizing the overall concordance in age association across the two technologies. (**B**) Predicted versus true ages for eight retinal cell classes in mouse Xenium datasets. Violin and box plots illustrate the distribution of predicted ages across chronological ages, with Pearson correlation coefficients (cor) provided in the upper left of each panel. (**C**) Scatter plots compare the aging clocks genes between Xenium and snRNA-seq for each cell type. Each point represents one aging clock gene. The x-axis represents the weights in snRNA-seq-based clocks and y-axis represents the weights in Xenium-based clocks. Most consistent aging clocks genes are labeled in the plot. The overall Pearson correlation coefficients between the two weights are indicated above each panel.

**Supplemental Figure 11: Examples of aging-regulated mouse genes showing concordant expression by snRNA-Seq and Xenium analysis.** (**A**) The boxplots showing examples of consistent aging related genes between Xenium and snRNA-seq datasets in Muller glia cells. The x axis indicate sages of the samples and y axis indicate z-score of the gene expression values.(**B**) Corresponding images of Xenium data for aging-related genes show in a. Scale bar: 500 μm.

## Supplemental Tables

**Supplemental Table 1: Cell Counts across aging in Zebrafish, Mouse, and Human.**

**Supplemental Table 2: List of all aging-regulated genes in zebrafish, mouse, and human.**

**Supplemental Table 3: GO and KEGG terms enriched in aging-regulated genes in zebrafish, mouse, and human.**

**Supplemental Table 4: Senescence-associated genes showing age-dependent regulation in major retinal cell types.**

**Supplemental Table 5: Common and distinct aging genes in the male and female human retina.**

**Supplemental Table 6: Genes used to calculate cell type-specific aging clocks.**

**Supplemental Table 7: Enriched and Conserved GO and KEGG Terms for Aging Clock Genes in zebrafish, mouse, and human.**

**Supplemental Table 8: List of aging-regulated DARs.**

**Supplemental Table 9: Enriched age-regulated transcription Factors and their overlap with age-dependent gene regulatory networks.**

**Supplemental Table 10: List of all aging-dependent cell-cells signaling interactions.**

**Supplemental Table 11: List of genes used to generate cell-specific aging clocks in Xenium dataset.**

