## Supplemental Figures S1-S11 for "Comparative single-cell multiomic analysis reveals evolutionarily conserved and species-specific cellular mechanisms mediating natural retinal aging"

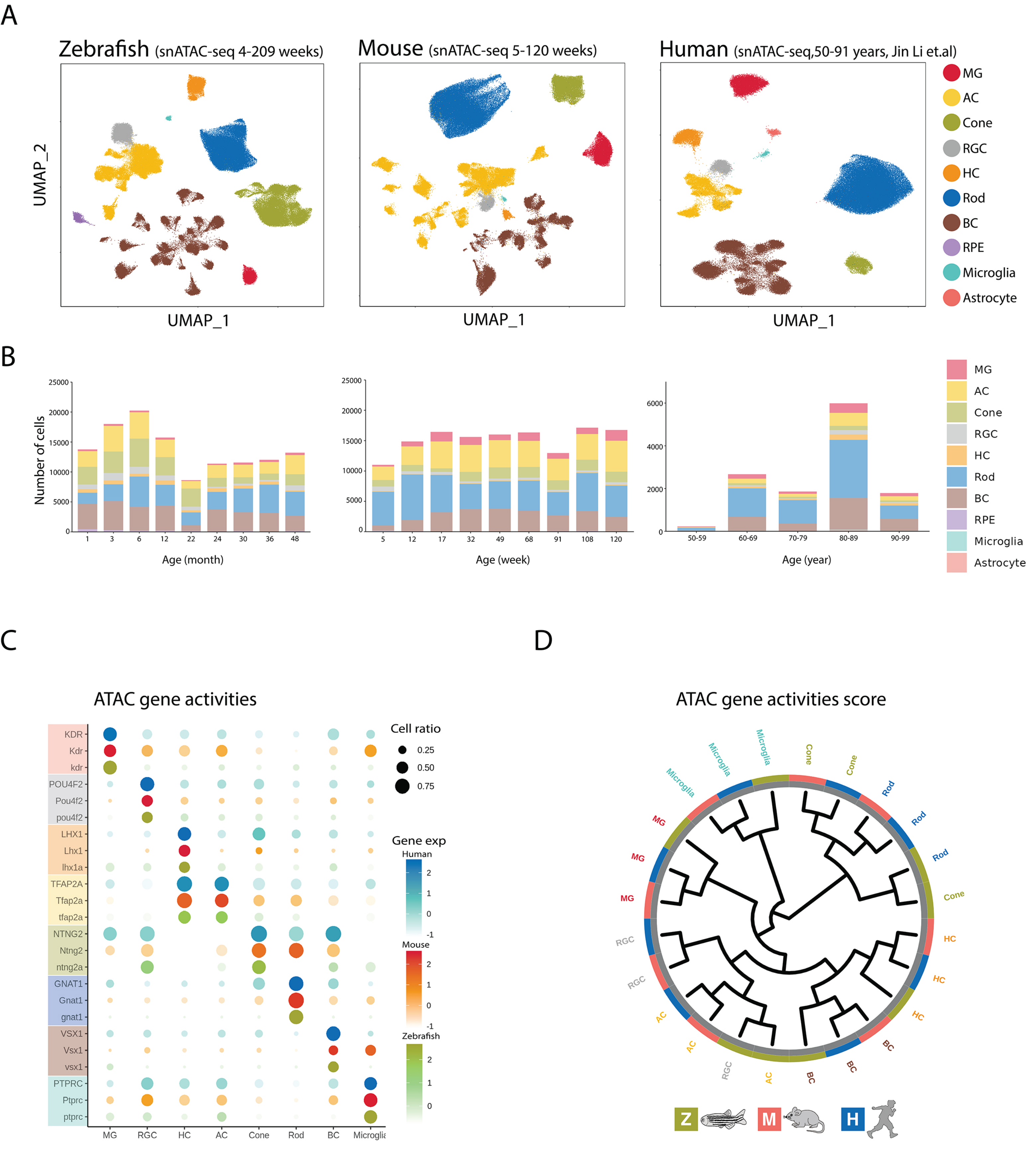


Lyu, et al. Fig. S1

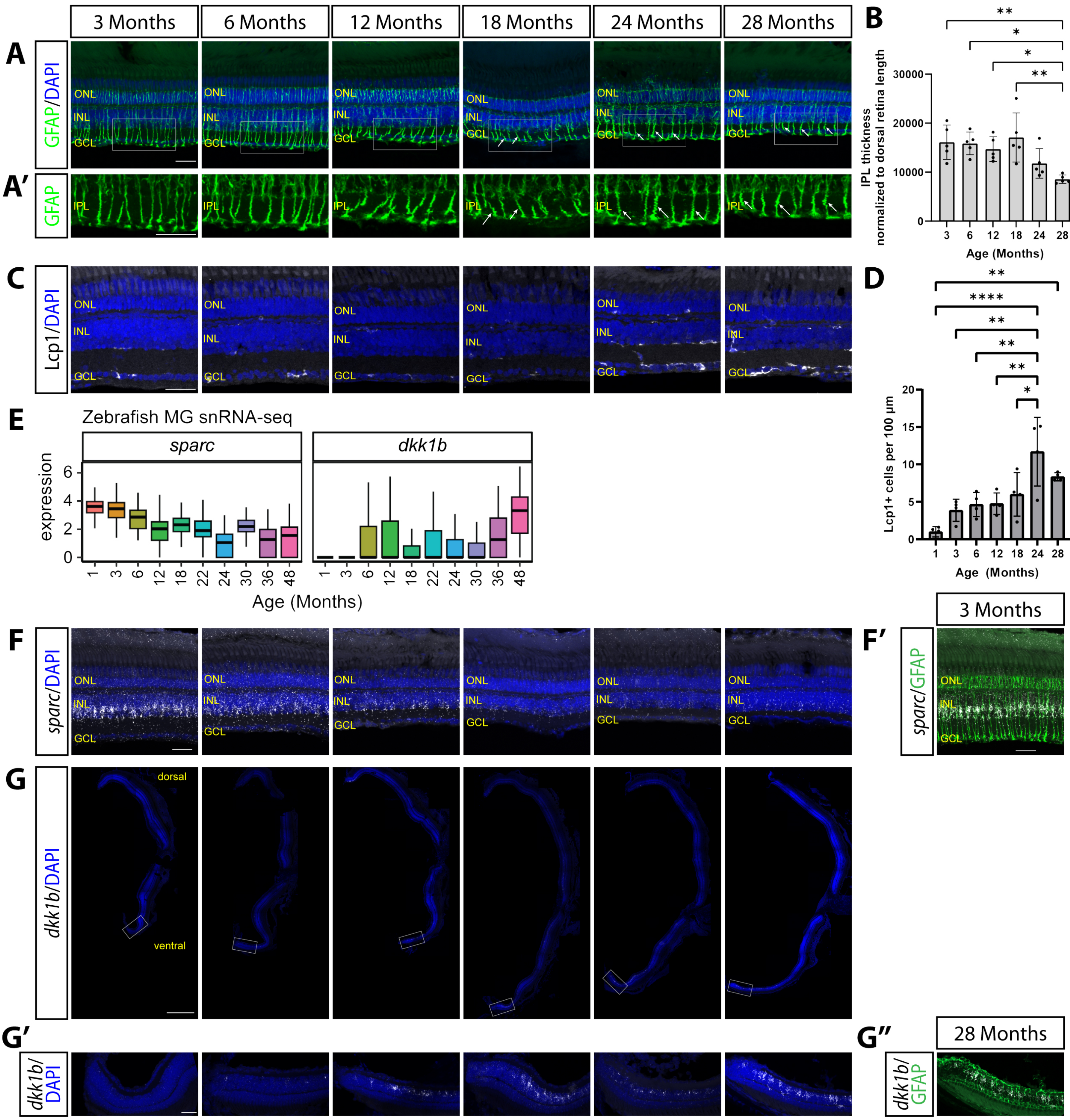


Lyu, et al Fig. S2

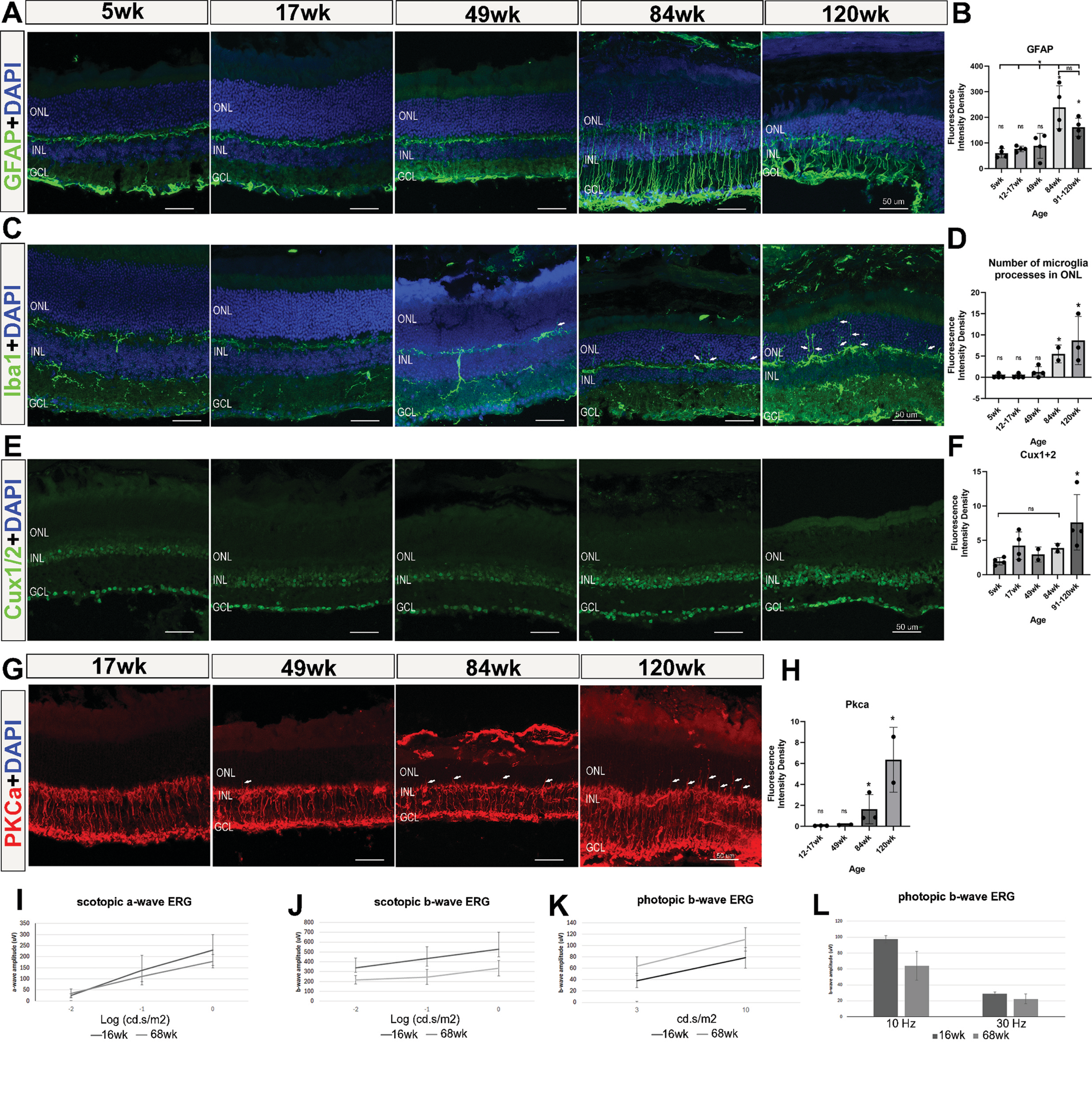


Lyu, et al. Fig. S3

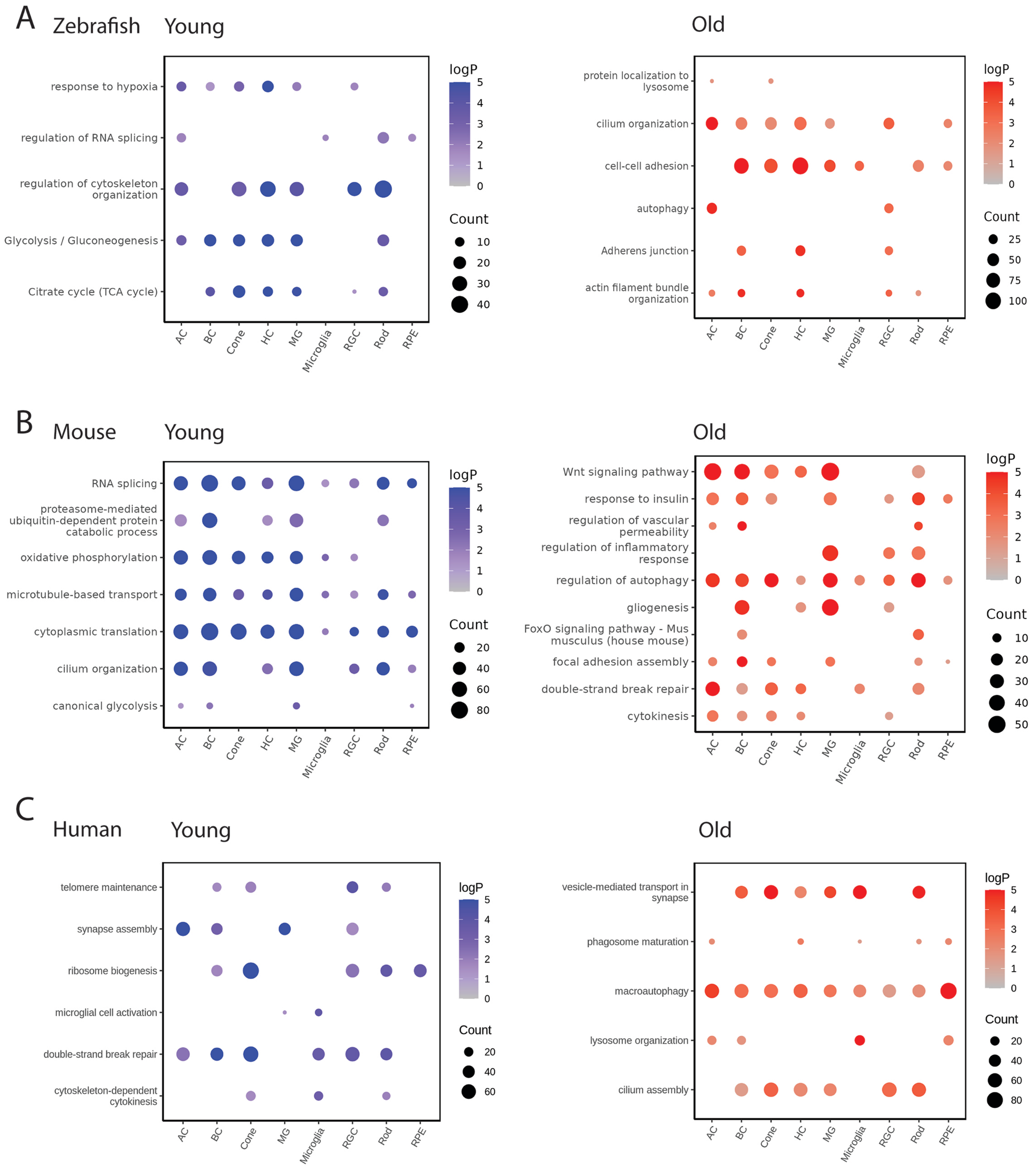


Lyu, et al. Fig. S4

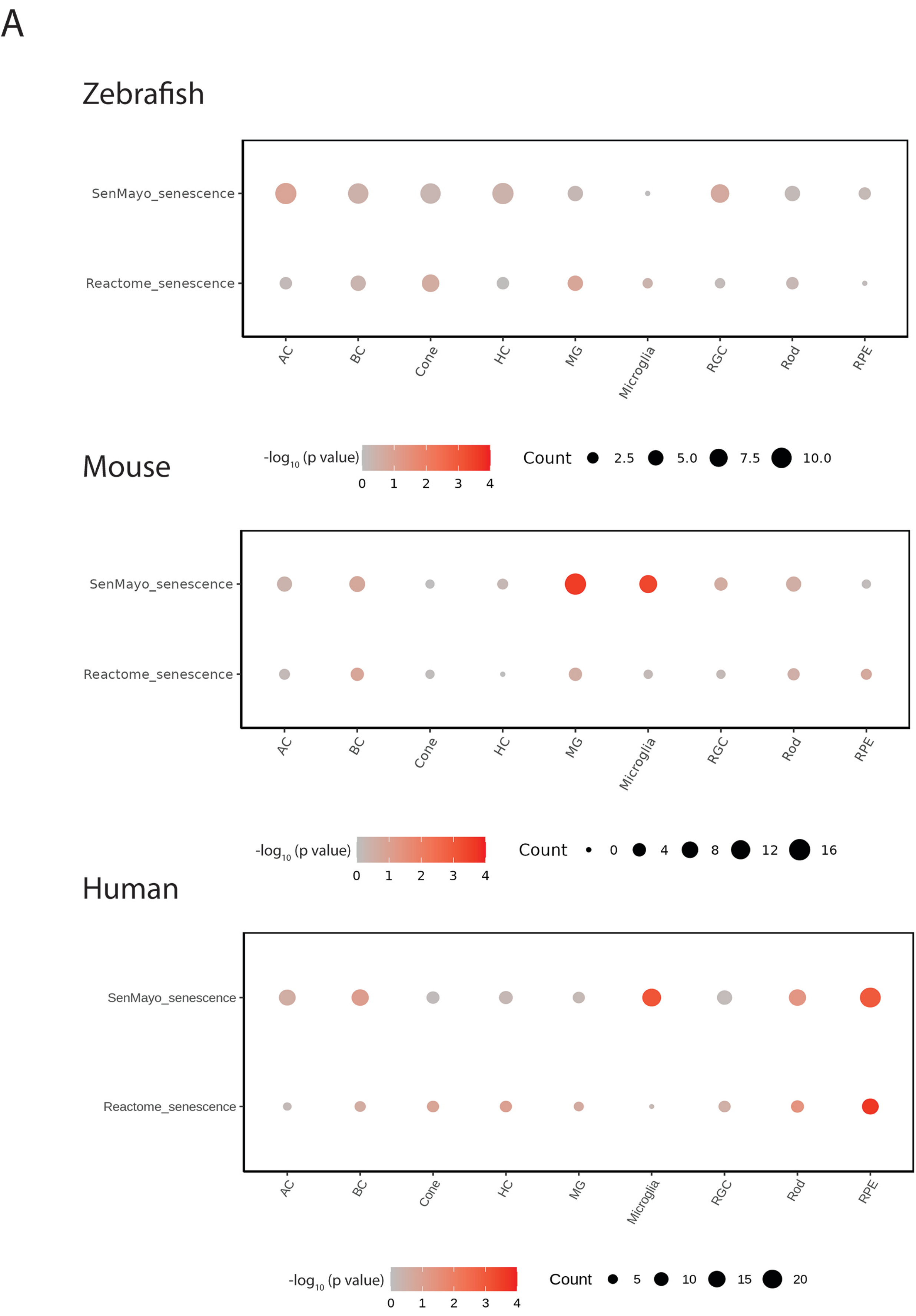


Lyu, et al. Fig.S5

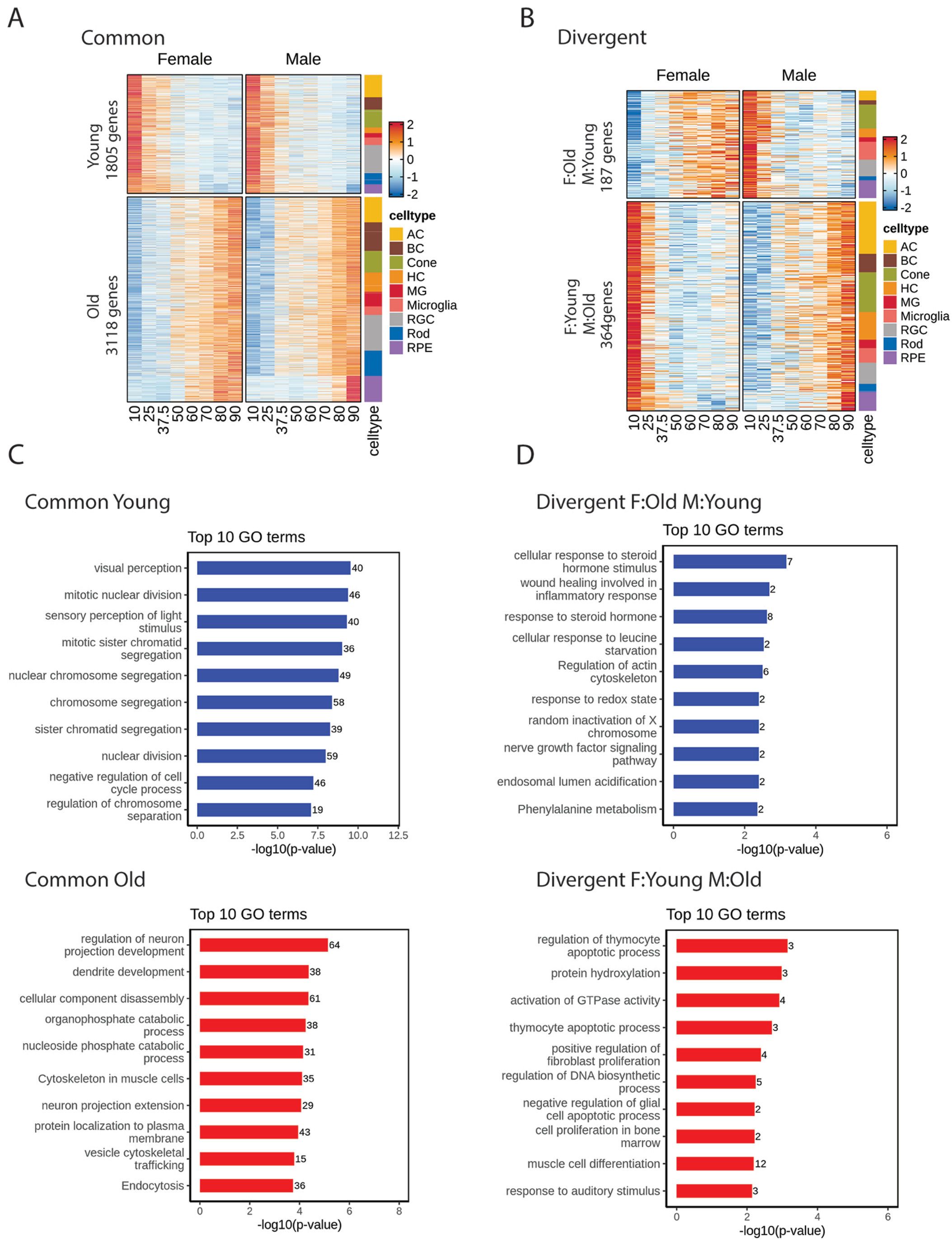


Lyu, et al. Fig.S6

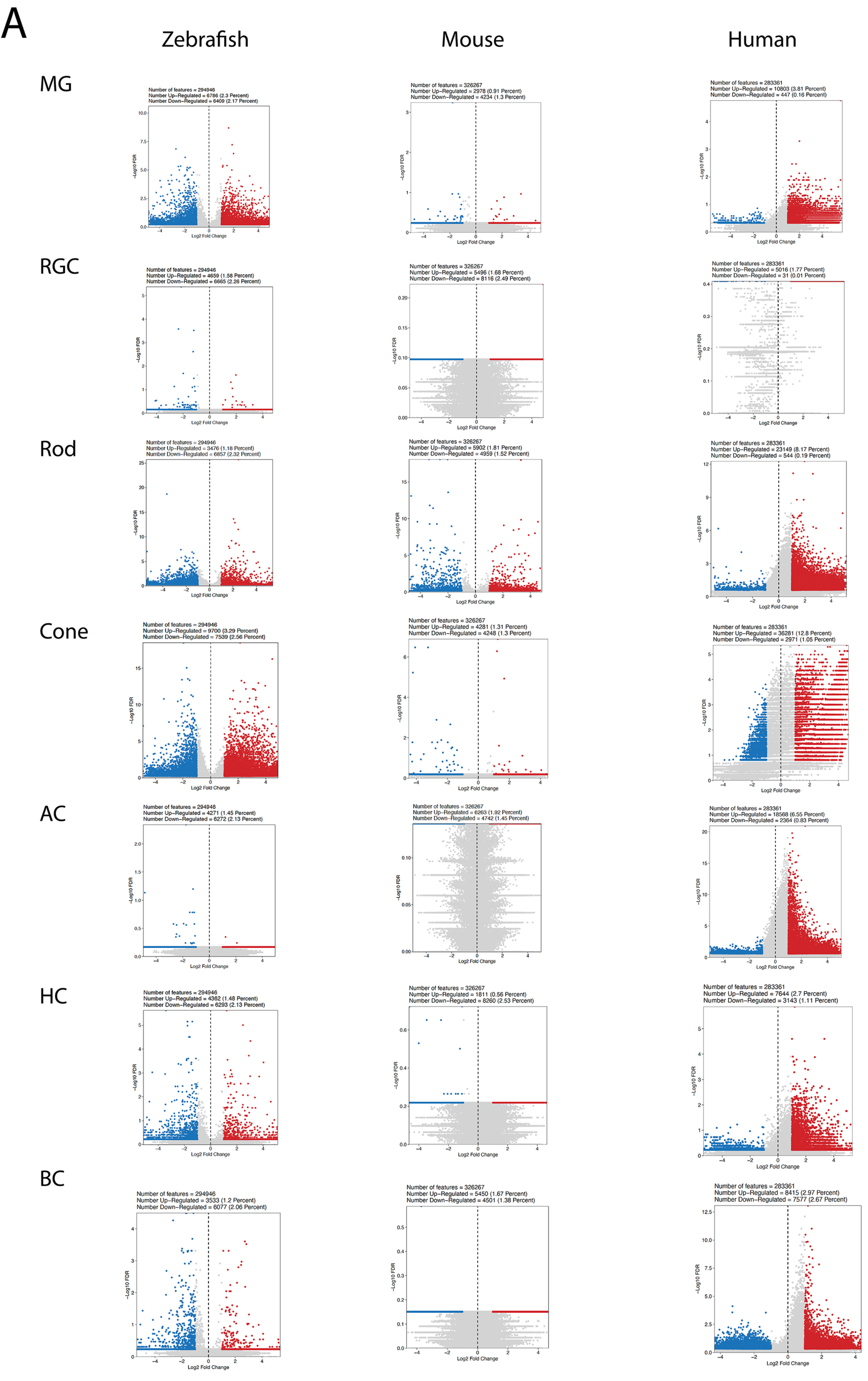


Lyu, et al. Fig. S7

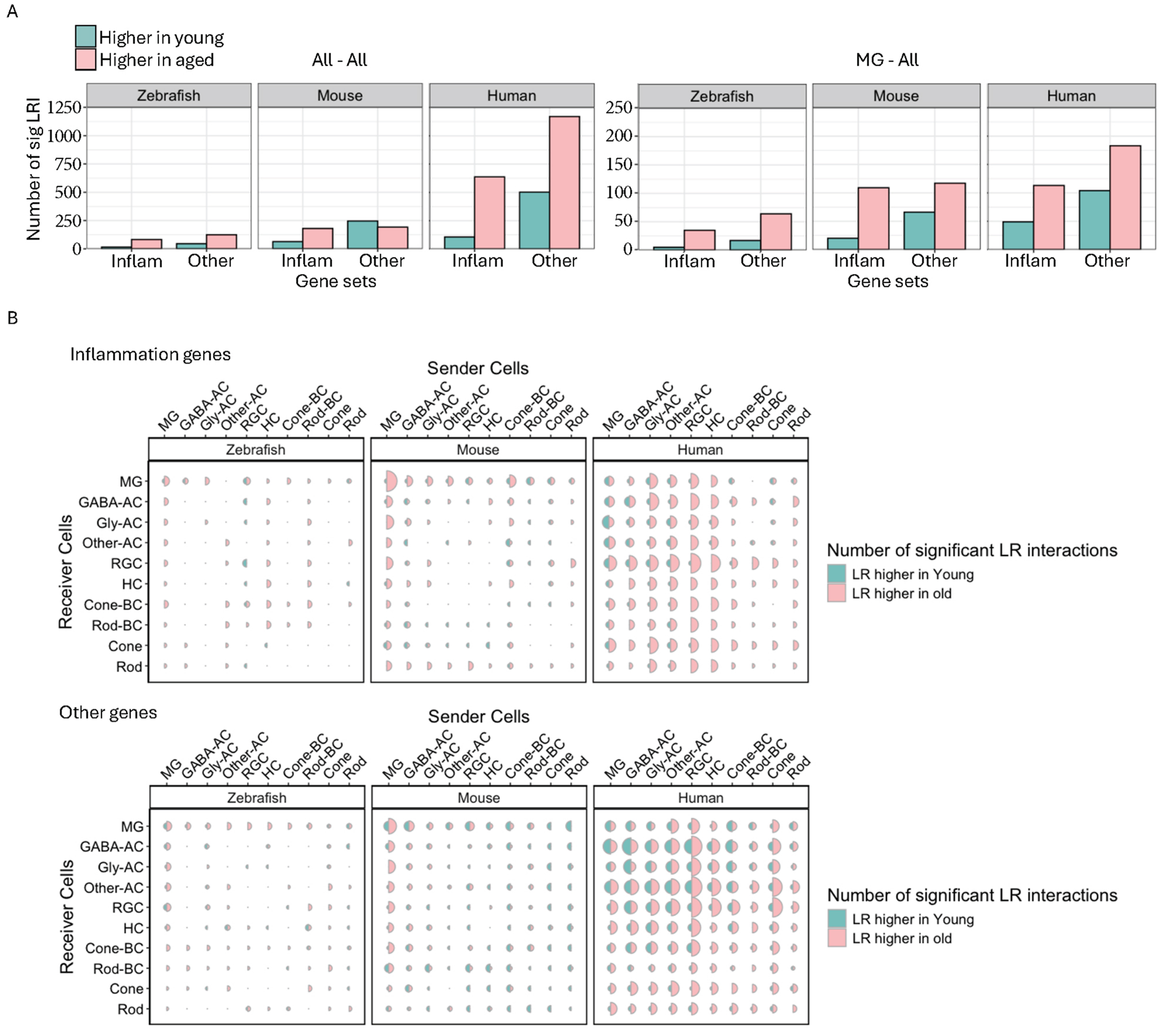


Lyu, et al. Fig. S8

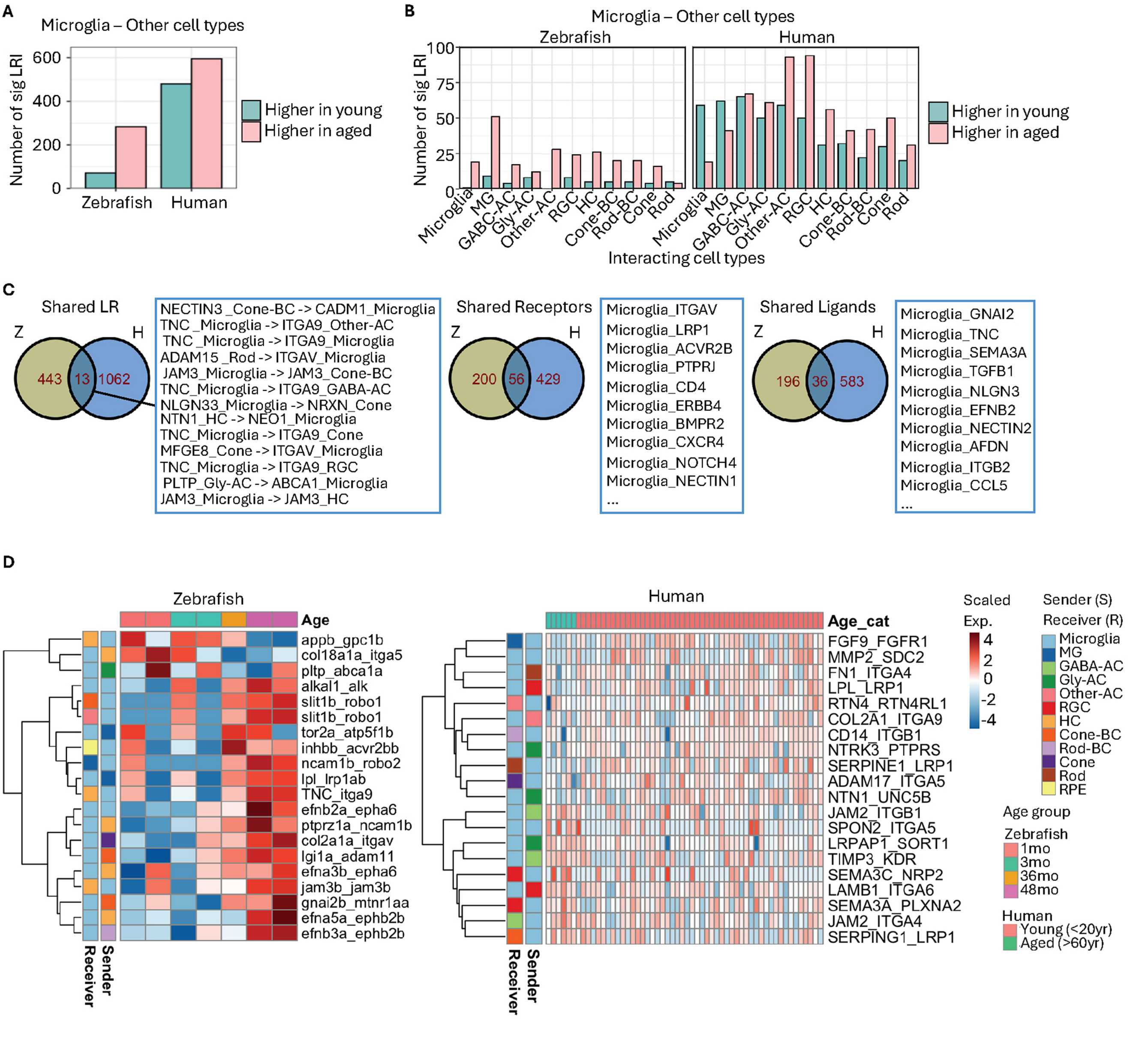


Lyu, et al. Fig. S9

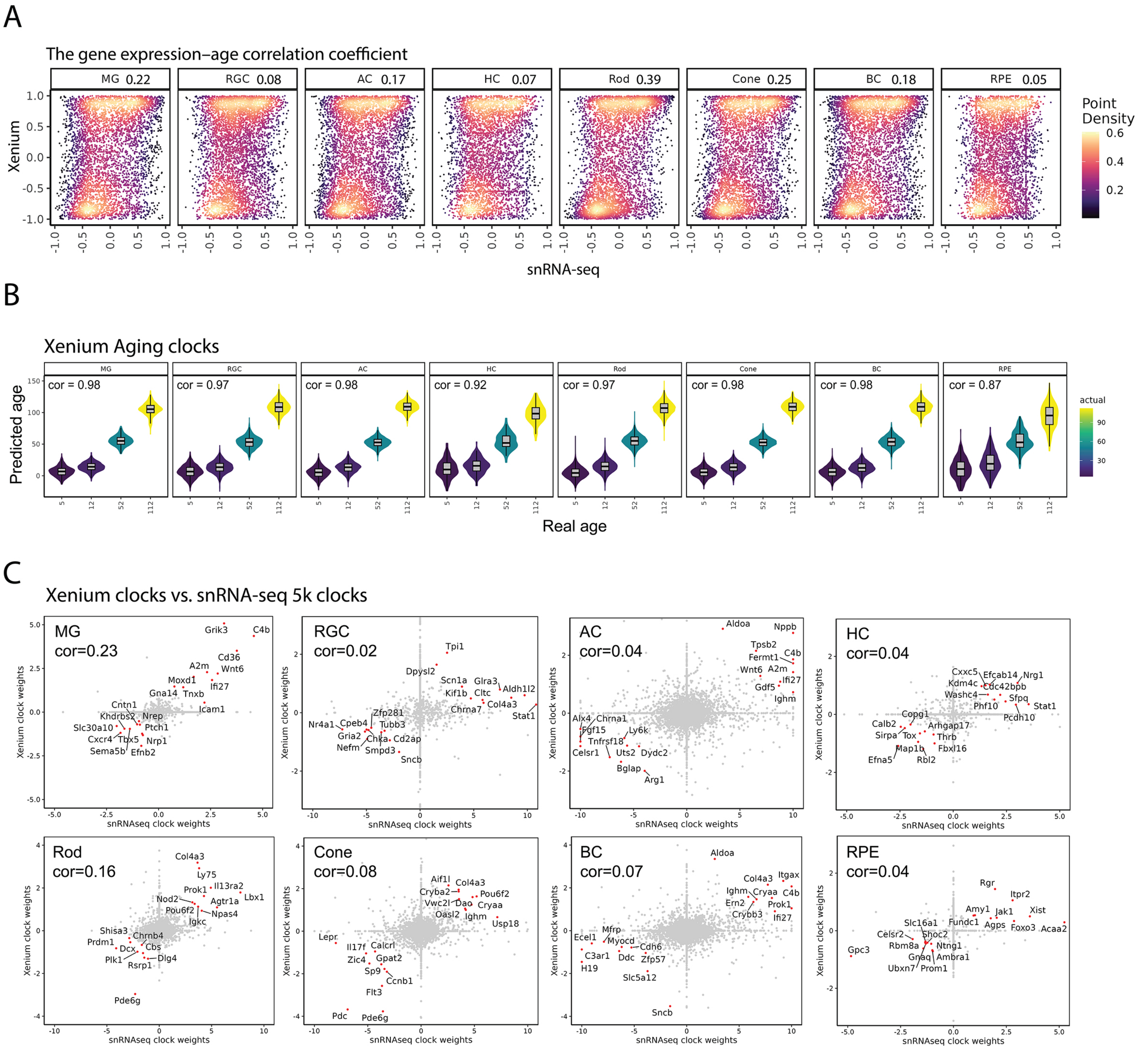


Lyu, et al. Fig. S10

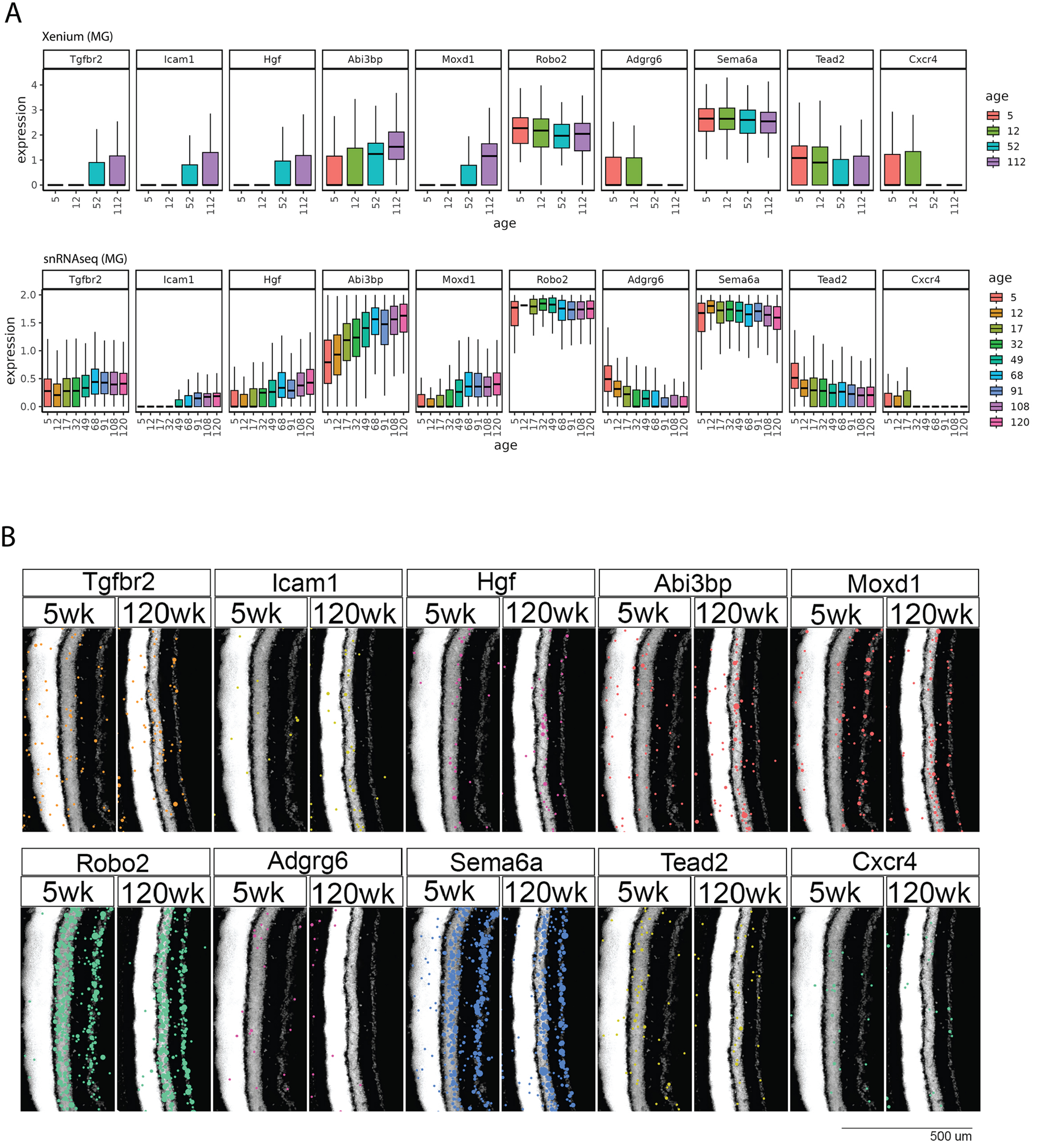


Lyu, et al. Fig. S11
